# Convergent utilization of APN receptor by hedgehog merbecoviruses

**DOI:** 10.1101/2025.09.16.676546

**Authors:** Peng Liu, Qianhui Zhu, Jingyi Liu, Zhixing Ma, Yuanling Yu, Chen Liu, Chengbao Ma, Junyu Si, Xiao Yang, Meiling Huang, Yehui Sun, Xiao Yu, Yucheng Sun, Tianci Liu, Lingling Yu, Qianran Wang, Jing Li, Yunlong Cao, Xiangxi Wang, Huan Yan

**Affiliations:** State Key Laboratory of Virology and Biosafety, Hubei Provincial Research Center for Basic Biological Sciences, College of Life Sciences, TaiKang Center for Life and Medical Sciences, Wuhan University; Wuhan, Hubei, China; Key Laboratory of Biomacromolecules (CAS), National Laboratory of Biomacromolecules, CAS Center for Excellence in Biomacromolecules, Institute of Biophysics, Chinese Academy of Sciences, Beijing, China; University of Chinese Academy of Sciences, Beijing, China; Biomedical Pioneering Innovation Center (BIOPIC), Peking University, Beijing, China; Changping Laboratory, Beijing, China; Peking-Tsinghua Center for Life Sciences, Tsinghua University, Beijing, China

**Keywords:** *Merbecovirus*, EriCoV, APN, receptor, hedgehog, TMPRSS2, Neutralizing antibodies

## Abstract

Dipeptidyl peptidase-4 (DPP4) and angiotensin-converting enzyme 2 (ACE2) are well-established receptors for merbecoviruses, yet the receptor usage of merbecoviruses of European and Asian hedgehogs (EriCoVs) remains unknown. Here, by testing hedgehog orthologs of known coronavirus receptors, we identify hedgehog aminopeptidase N (APN) as a functional receptor for EriCoVs. Analysis of 94 APN orthologs indicates that EriCoVs have a limited host range, primarily utilizing hedgehog APN and, to a lesser extent, APN from certain felids, shaped by specific determinants at the virus–receptor interface. Cryo-EM reveals an APN-binding mode distinct from those used by alpha- and deltacoronaviruses. Functional assays indicate that hedgehog transmembrane serine protease 2 (TMPRSS2) enhances spike activation and promotes pseudovirus entry. Neutralizing antibodies targeting RBD and APN were developed and could effectively block EriCoV pseudovirus entry and propagation. These findings reveal an unexpected convergent evolution of APN utilization among merbecovirus, establishing a foundation for risk assessment and countermeasure development.

**Graphic abstract:** Convergent utilization of hedgehog APN by EriCoVs

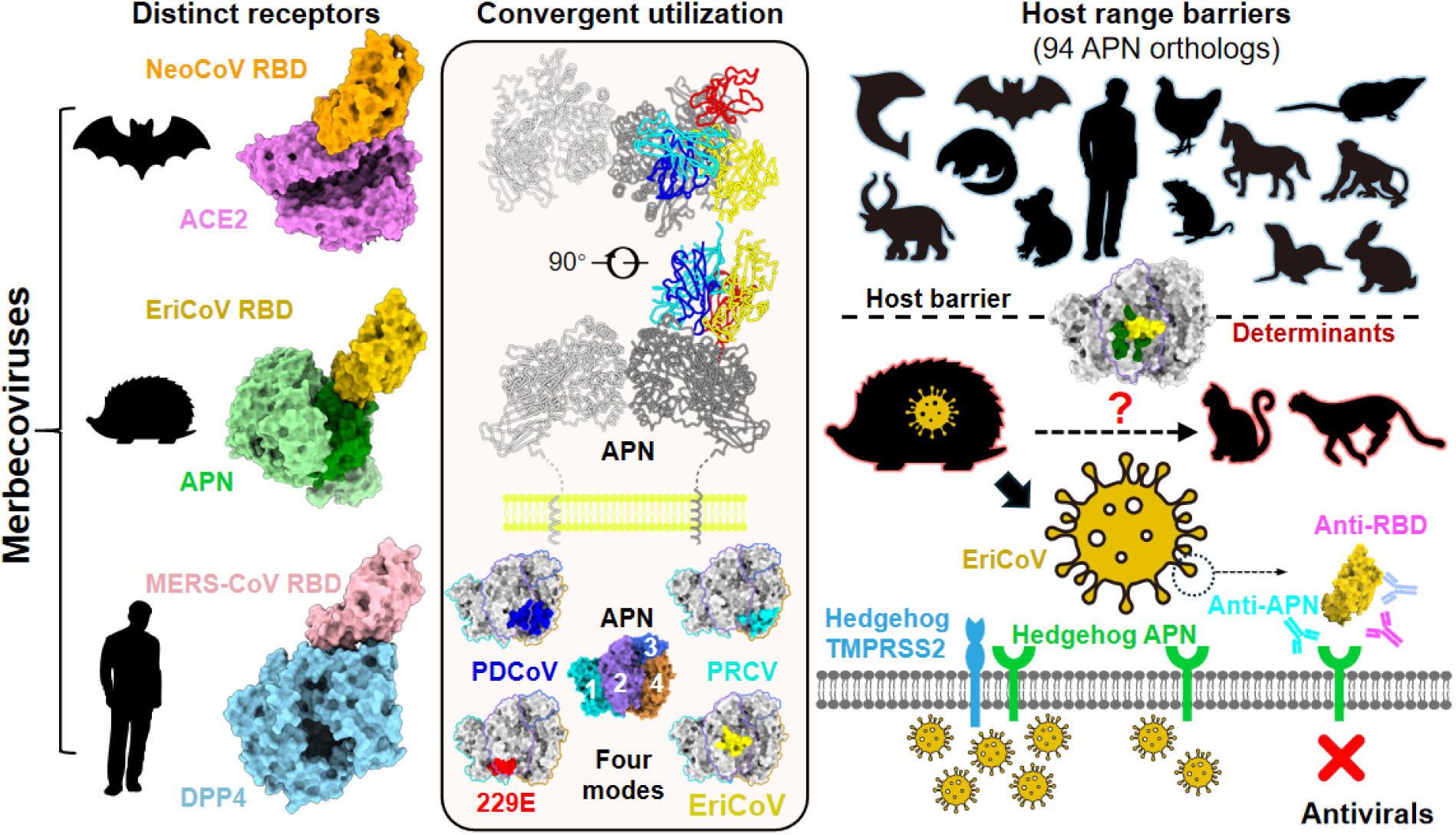

## INTRODUCTION

Three human betacoronaviruses—SARS-CoV-1, MERS-CoV, and SARS-CoV-2—have caused devastating outbreaks in the 21^st^ century.^1–5^ Among betacoronaviruses, the *Merbecovirus* subgenus has been designated a priority pathogen group by the World Health Organization (WHO).^6,7^ These viruses include the highly pathogenic Middle East respiratory syndrome coronavirus (MERS-CoV), which infects both humans and camels,^3,8,9^ along with numerous relatives with varying degrees of zoonotic potential.^10–12^

Among them, *Betacoronavirus cameli* (MERSr-CoV), *Betacoronavirus pipistrelli* (HKU5), and *Betacoronavirus tylonycteridis* (HKU4) are three viral species that originated in bats^13–18^ and have spilled over into mammals such as humans,^3^ camels,^19–21^ pangolins,^22,23^ and minks.^24^ In 2013, a novel merbecovirus species (*Betacoronavirus erinacei*, or EriCoV/ErinCoV) was discovered in fecal samples from European hedgehogs (*Erinaceus europaeus*, *E.eur*) in Germany.^25^ A related Asian lineage (HKU31) was later found in Amur hedgehogs (*Erinaceus amurensis*, *E.amu*) in China.^26^ To date, EriCoVs have been reported exclusively in hedgehogs across a broad geographic range across Euroasia, including Germany,^25^ France,^27^ United Kingdom,^28^ Italy,^29,30^ Poland,^31,32^ Portugal,^33^ Russia,^34^ and China.^26,35,36^ Among them, all Asian EriCoVs were sampled in *E.amu* hedgehogs, while nearly all European EriCoVs were found in *E.eur* hedgehogs, except for two strains from Moscow annotated as *Erinaceus sp.*.^34^ Despite their broad geographic distribution and genetic diversity, EriCoVs remain poorly understood in terms of pathogenicity and cross-species potential.^27,33^

The host range and transmissibility of coronaviruses are primarily determined by interactions between the spike (S) protein receptor-binding domain (RBD) and specific cellular receptors.^37^ So far, at least five host proteins have been established as functional coronavirus receptors: carcinoembryonic antigen-related cell adhesion molecule 1a (CEACAM1a),^38,39^ angiotensin-converting enzyme 2 (ACE2),^40–42^ dipeptidyl peptidase-4 (DPP4),^14,15,43^ aminopeptidase N (APN),^44,45^ and transmembrane protease serine 2 (TMPRSS2).^46–49^ Among them, ACE2 and APN are recognized by phylogenetically diverse coronaviruses with distinct RBD structures, indicating multiple independent receptor acquisition events during evolution.^50–52^ ACE2 usage is shared by coronaviruses across alpha- and betacoronavirus, including setracoviruses (e.g., NL63^41^), sarbecoviruses (e.g., SARS-CoV-2^5,53,54^), and several merbecovirus clades (e.g., NeoCoV,^40^ MOW15-22,^51^ HKU5,^24,50,55–57^ HKU25^58^). APN serves as a receptor for multiple alpha- and deltacoronaviruses, including duvinacoviruses (e.g., 229E^44^), tegacoviruses (e.g., TGEV,^45^ CCoV-HuPn-2018,^59^ FCoV-23^60^), and buldecoviruses (e.g., PDCoV,^61^ and multiple Avian deltacoronaviruses^62^). TMPRSS2, besides serving as a functional receptor for HKU1, is also a critical protease for spike activation of many coronaviruses, influencing the route and tropism of these viruses.^54,63–65^

While DPP4 has been considered the canonical receptor for merbecoviruses, recent findings suggest that ACE2 may be more prevalent than previously apprehended.^40,50,51,57^ In 2022, we reported that NeoCoV and PDF-2180, close relatives of MERS-CoV in African bats, use ACE2 as their functional receptors.^40,66^ Subsequently, several other merbecovirus clades were also found to acquire ACE2 usage independently.^50,51,57,58^ However, EriCoVs do not use either ACE2 or DPP4 for entry, leaving their entry and transmission mechanism unresolved.^40,56,67^

Here, we show that EriCoVs use hedgehog APN as an entry receptor and rely on hedgehog TMPRSS2 for spike activation. Cryo-EM analyses unveil a previously unrecognized APN-binding mode, representing another unique case of convergent APN utilization by betacoronavirus. This study provided the last major piece to the long-standing puzzle of merbecovirus receptor usage, and points the way toward effective assessment and mitigation of the zoonotic threats posed by those MERS-CoV relatives.

## RESULTS

### Characterization of EriCoV receptor binding domain

To investigate the receptor binding features of EriCoVs, we analyzed 24 publicly available EriCoV genomic sequences and 25 EriCoV S glycoprotein sequences, with particular focus on the RBD regions. For conciseness and clarity, EriCoV strain names were abbreviated in this study (Figure S1A and Data S1). Among these, 5 Asian EriCoV strains (HKU31) were identified in Amur hedgehogs (*E. amurensis*) from several provinces of China, while 20 European EriCoV strains were identified in various European countries, predominantly confirmed from European hedgehogs (*E. europaeus*) (Figure 1A).

**Figure 1.**
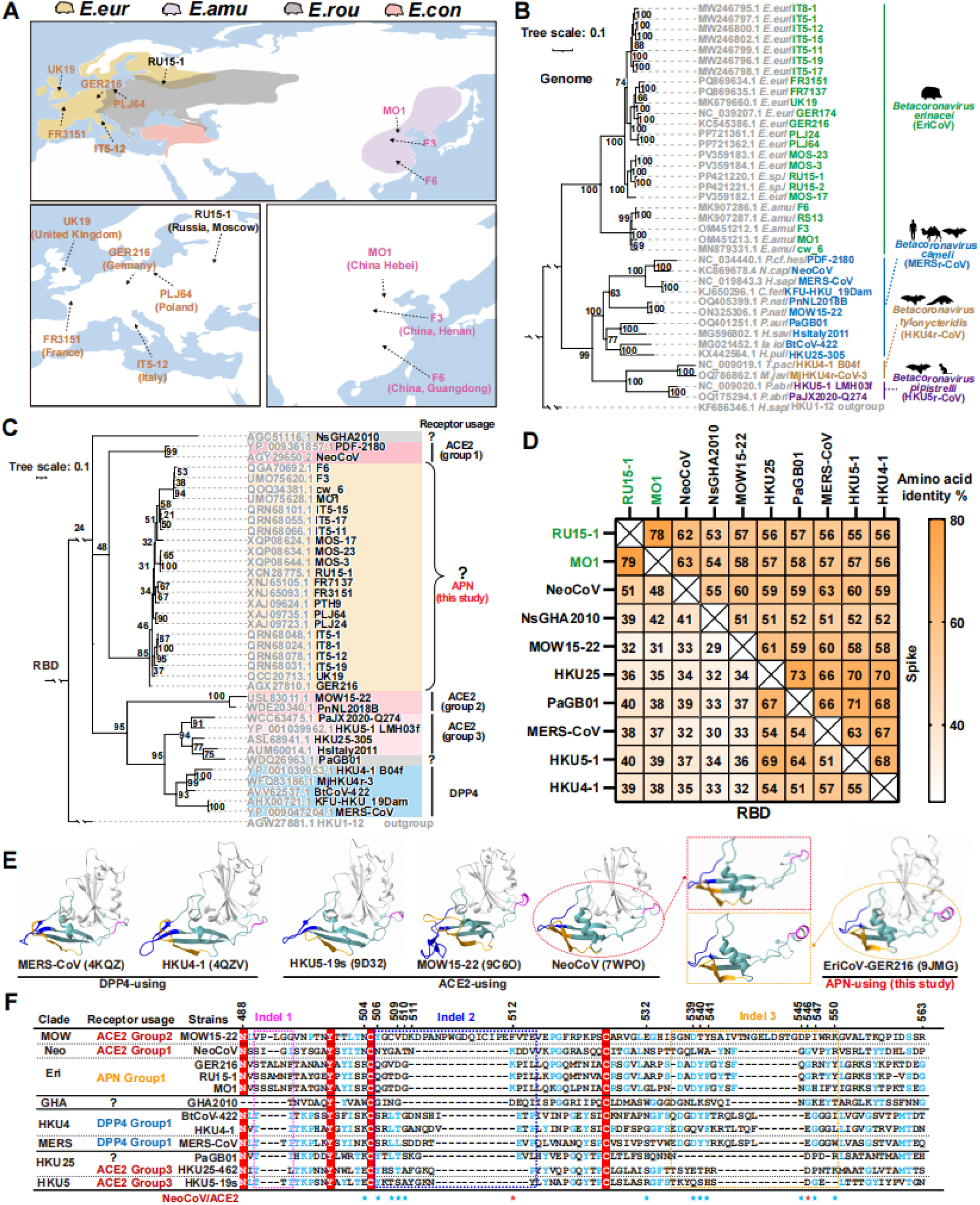
Geographic distribution, phylogenetic relationship, and RBD features of EriCoVs. **(A)** Geographic distribution of Erinaceus hedgehogs and reported sampling locations of selective EriCoV strains. Host abbreviations: *E.eur* (*Erinaceus europaeus*), *E.amu* (*Erinaceus amurensis*), *E.rou* (*Erinaceus roumanicus*), *E.con* (*Erinaceus concolor*). Species distribution data retrieved from the International Union for Conservation of Nature (IUCN) Red List were mapped using GeoScene Pro. The color scheme of EriCoVs corresponds to their host, except for RU15-1 with an unconfirmed hedgehog species (*Erinaceus sp.*). (**B-C**) Maximum-likelihood phylogenetic trees of representative merbecoviruses based on complete genome nucleotide sequences (B) or RBD amino acid sequences (corresponding to NeoCoV residues 380–585) (C) were generated by IQ-TREE. HKU1-12 was designated as the outgroup. Host, viral species, receptor usage, and receptor binding mode are annotated. ACE2-using merbecoviruses were grouped according to evolutionarily distinct receptor-binding modes characteristic of the NeoCoV-(clade 1), MOW15- 22- (clade 2), and HKU5/HKU25-related (clade 3) clades. Scale bars denote genetic distance (substitutions per site). (**D**) Pairwise amino acid identity matrix of receptor-binding domains (RBDs) and full-length S glycoproteins for selected merbecoviruses. EriCoV strains were labeled in green. (**E**) Structures of representative merbecovirus RBDs. The conserved RBD core subdomains are rendered in gray, and receptor-binding motifs (RBMs) are rendered in dusty teal; structurally divergent RBM indels are highlighted (magenta: indel 1; dark blue: indel 2; orange: indel 3). The magnified RBMs from NeoCoV and EriCoV (GER216) were shown to facilitate comparison. (**F**) Manually adjusted multiple sequence alignment of RBMs with optimized indel positioning. Fully conserved residues are shown with a red background; partially conserved residues are colored with light blue. Dashed boxes define indels that correspond with panel E. Residues mediating ACE2 interactions by NeoCoV are asterisked (red: conserved in EriCoVs; blue: divergent). NeoCoV residue numbering is shown. See also Figure S1.

Phylogenetic analyses of both genomic sequences and S glycoprotein amino acid sequences revealed that Asian and European EriCoVs form sister lineages, distant from bat-borne merbecoviruses (Figures 1B and S1B). This separation suggests a long-term evolutionary divergence of EriCoVs in hedgehogs across Eurasia. Consistent with previous findings, EriCoV RBD sequences cluster with those of ACE2-using NeoCoV and PDF-2180, implying a potential shared ancestry (Figure 1C).^40^ However, due to limited sequence homology and likely different receptor usage patterns, we classified EriCoVs and NeoCoV-related viruses into distinct clades in this study. Among the analyzed sequences, pairwise amino acid sequence analysis of S and RBD shows that two representative EriCoV strains from Asian and European, respectively, share moderate sequence identity (78∼79%), but both are markedly divergent from other merbecoviruses (32∼51% for RBD and 53∼63% for S), with NeoCoV being the closest known strain (Figure 1D).

Structural comparison of RBDs from representative merbecoviruses revealed that EriCoV and NeoCoV share a similar overall RBD core and receptor-binding motif (RBM) architecture, including characteristic insertion–deletion (indel) patterns (Figure 1E), which have been implicated in receptor usage shifts during coronavirus evolution.^58,68^ Further examination of the putative RBM sequences confirmed shared indel patterns between NeoCoV and EriCoVs, except for indel 1, which lies outside the canonical interaction interface. However, only 2 out of 14 ACE2-interacting NeoCoV residues are conserved in EriCoV,^40^ suggesting a distinct receptor engagement mechanism (Figure 1F).

Collectively, these analyses suggest that EriCoVs likely employ a novel receptor-binding strategy that does not involve known ACE2 or DPP4 interaction modes, concurring with the observations in prior studies.^40,56^

### Hedgehog APN is a receptor for EriCoV

Given previous failed attempts to identify receptor functionality among various mammalian ACE2 orthologs^40^ and the hypothesis that EriCoVs may utilize an uncharacterized receptor binding mode, we explored whether these viruses engage alternative known coronavirus receptors from hedgehog hosts, including DPP4, APN, and the recently identified HKU1 receptor TMPRSS2. To this end, we retrieved protein-coding sequences of ACE2, DPP4, TMPRSS2, and APN from the RNA-seq data of *E. amurensis* hedgehog (Genome Sequence Archive, GSA database)^69^ for subsequent evaluation. Notably, APN amino acid sequences were identical between *E.amu* and *E.eur* hedgehogs, allowing *E.amu* APN to serve as a representative for both hedgehog host species (Data S2).

We first tested two representative EriCoV strains: RU15-1 (European) and MO1 (Asian). Among the tested receptors, only *E.amu* APN, but not *E.amu* ACE2, DPP4, TMPRSS2, or human (h) counterparts, supported detectable RBD binding (Figure 2A) and RU15-1 and MO1 pseudovirus entry (Figures 2B and 2C). Flow cytometry confirmed specific binding of EriCoV RBDs to *E.amu* APN but not to hAPN (Figure 2D). A pull-down assay further demonstrated that EriCoV RBD, but not 229E RBD, interacted specifically with *E.amu* APN (Figure 2E). Notably, mutation of the APN catalytic site^70^ did not impair IT5-12 RBD binding or pseudovirus entry, indicating that receptor function is independent of protease activity (Figures S2A and S2B). Although hAPN was not an efficient receptor for EriCoVs, CRISPR-Cas9 knockout (KO) of endogenously expressed hAPN abolished the limited EriCoV S-mediated membrane fusion (Figure 2F) and pseudovirus entry in Caco-2 cells (Figure 2G), reducing both to background levels. A similar phenotype was observed for 229E, which was used as a positive control (Figure 2G).

**Figure 2.**
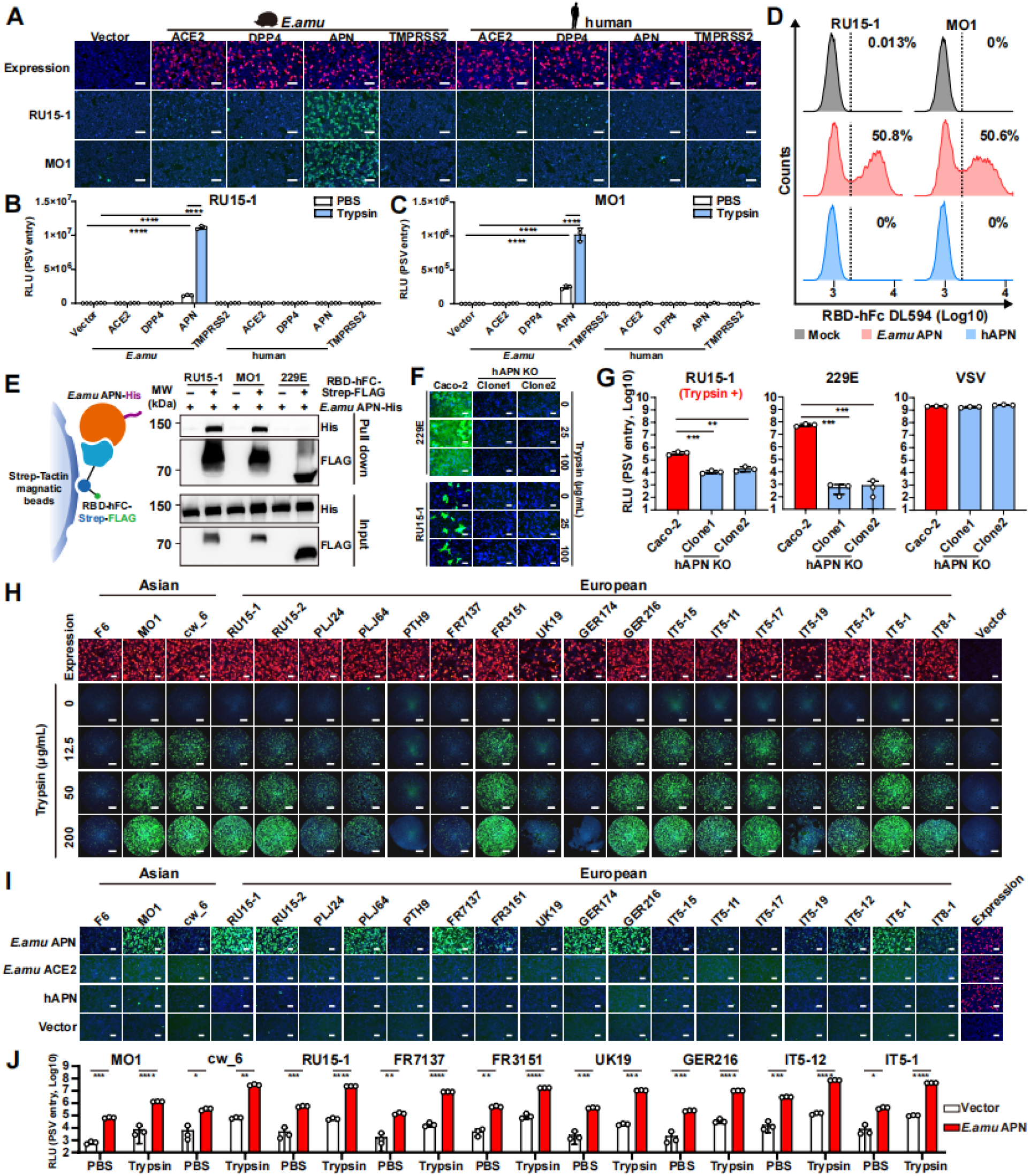
Hedgehog APN functions as a receptor for EriCoVs. (**A-C**) Transient expression of *E. amurensis* (*E.amu*) APN, but not *E.amu* ACE2, DPP4, TMPRSS2, nor any tested human receptor counterparts, supported EriCoV RU15-1 and MO1 RBD human IgG1 Fc domain (hFc) fusion protein (RBD-hFc) binding (A), as well as pseudovirus entry (B-C) in HEK293T cells. Receptor expression was confirmed by immunofluorescence staining: FLAG-tagged ACE2, DPP4, and APN, and ALFA-tagged TMPRSS2 (red). Pseudoviruses (PSV) were pretreated with 1.25 mg/mL trypsin. (**D**) Flow cytometry analyses of RU15-1 and MO1 RBD-hFc binding to HEK293T cells transiently expressing hAPN or *E.amu* APN. Dashed lines: background threshold. Means of three technical repeats are shown. (**E**) Pull-down assay showing specific interaction of soluble *E.amu* APN ectodomain (His-tagged) with Strep-Tactin–immobilized RU15-1 or MO1 RBD-hFc-Strep-FLAG fusion proteins, with 229E RBD as a negative control. (**F**) Cell–cell fusion assays in Caco-2 or hAPN-knockout (KO) Caco-2 cells (two clones) transiently expressing 229E or RU15-1 spike proteins, with or without trypsin treatment. (**G**) Infection of Caco-2 and hAPN-KO Caco-2 cells by the indicated pseudoviruses. RU15-1 pseudoviruses were pretreated with 1.25 mg/mL trypsin, and infection was performed in serum-free DMEM containing 125 μg/mL trypsin. (**H-I**) S-mediated cell-cell membrane fusion (**H**) and S_1_-hFc binding (**I**) of indicated Asian and European EriCoV strains in HEK293T cells stably expressing *E.amu* APN with the treatment of indicated concentrations of trypsin for panel H, or in HEK293T cells transiently expressing *E.amu* APN, *E.amu* ACE2, hAPN, or vector control for panel I. S glycoprotein and receptor expression was evaluated by immunofluorescence, detecting the C-terminally fused HA tag in panel H, and C-terminally fused FLAG tag in panel I (red). (**J**) Trypsin-pretreated (1.25 mg/mL trypsin) pseudovirus entry efficiency of selected EriCoV strains in HEK293T cells transiently expressing the *E.amu* APN. Mean ± SD and n=3 biological replicates for B, C, G, and J. Statistical significance was determined using unpaired two-tailed *t* tests. **: *P* < 0.01, ***: *P* < 0.005, and ****: *P* < 0.001. RLU: Relative light unit. Scale bars: 100 μm for A, F, I, and 5 mm for H. See also Figure S2.

We next evaluated whether *E.amu* APN also serves as a functional receptor for additional EriCoV strains. In a cell-cell fusion assay, *E.amu* APN promoted spike-mediated membrane fusion for most tested strains in a trypsin-dependent manner (Figures 2H and S2C). Similarly, *E.amu* APN, but not *E.amu* ACE2 or hAPN, supported S_1_-hFc and RBD-hFc binding from multiple EriCoV strains to varying efficiencies (Figure 2I and S2D). These findings were further validated by entry assays using VSV pseudotyped with S glycoproteins from diverse EriCoV strains (Figures 2J and S2E).

Together, these data demonstrate that APN, but not other known coronavirus receptors, serves as a functional receptor for both Asian and European EriCoVs through specific interaction with their RBDs.

### Ortholog specificity and host tropism determinants

To assess the potential host range and spillover risk of EriCoVs, we examined the ability of various APN orthologs to support RBD binding and pseudovirus entry when transiently expressed in HEK293T cells. We established an APN library comprising 94 orthologs spanning 13 taxonomic orders (31 bats, 62 non-bat mammalian species, and one avian species), all with validated expressions (Figures 3A, S3A, and Data S3).

**Figure 3.**
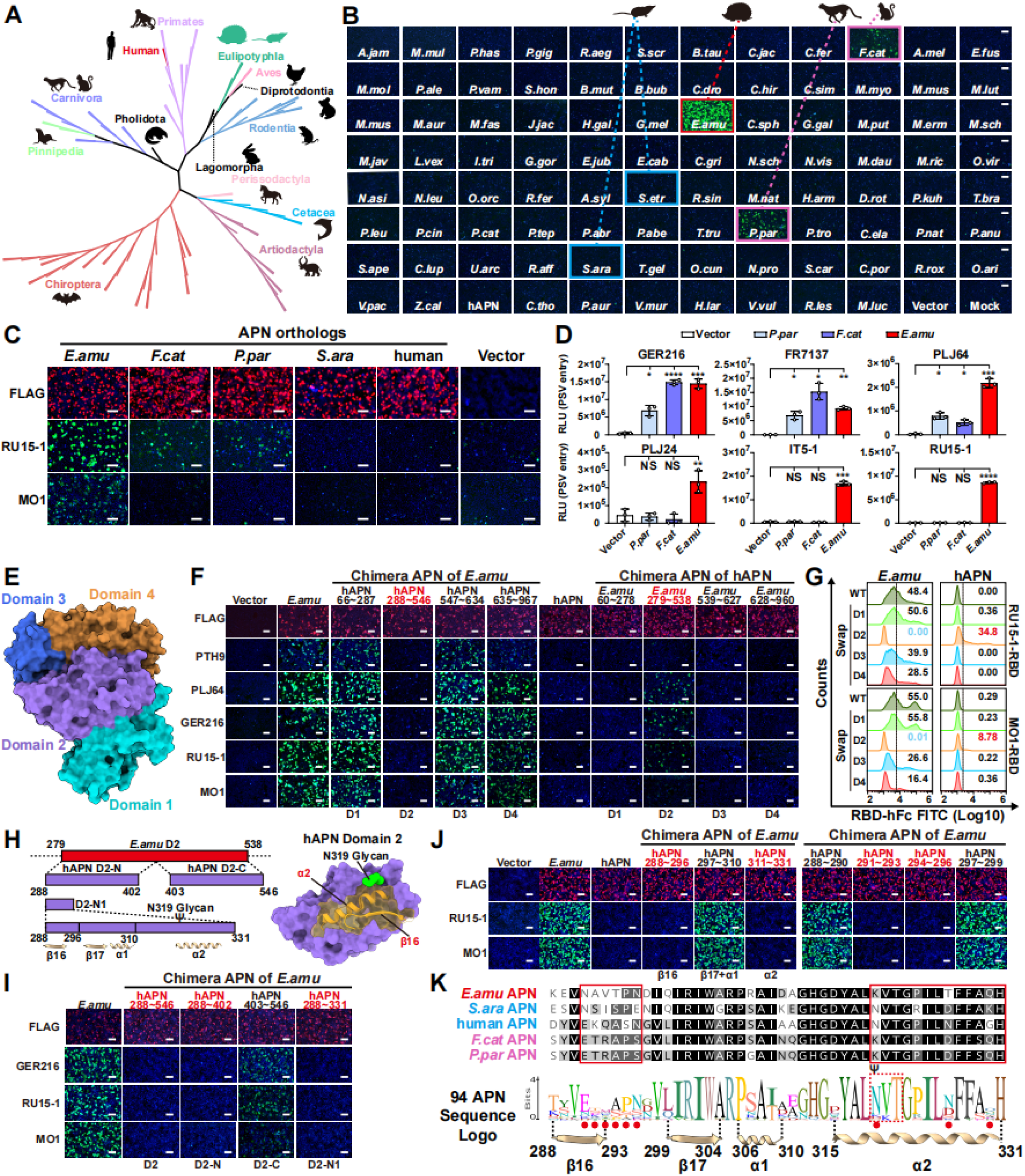
Multi-species APN tropism and host range determination of EriCoV. (**A**) Maximal likelihood phylogenetic trees of 94 APN orthologs from mammalian and avian species were constructed using IQ-TREE based on amino acid sequences. Taxonomic orders were annotated. Ortholog details are provided in Data S3. (**B**) Binding of RU15-1 S_1_-hFc (green) to HEK293T cells transiently expressing the indicated APN orthologs, assessed by immunofluorescence using an anti-hFc antibody. Red box: efficient binding. Magenta box: detectable binding. Blue box: two Eulipotyphla species tested with undetected binding. (**C**) RU15-1 and MO1 RBD-hFc binding (green) to HEK293T cells transiently expressing the indicated APN orthologs. (**D**) Entry of selected EriCoV strains into HEK293T cells transiently expressing indicated APN orthologs. PSVs were pretreated with 1.25 mg/mL trypsin. (**E**) Structural model of human APN with four domains highlighted in different colors. (**F-G**) Immunofluorescence (F) and flow cytometry (G) analyses of the indicated EriCoV RBD-hFc binding to HEK293T cells transiently expressing APN chimeras with domain swaps between human and *E.amu* APNs. Dashed lines: background threshold. Data shown are the mean of three technical repeats. (**H**) Schematic illustration of *E.amu* APN chimeras with equivalent residues from hAPN. Secondary structures and their surface localization of β16 and α2 in domain 2 (D2) were illustrated. (**I-J**) Fine mapping of the host range determinant restricting hAPN binding with RBD-hFc from RU15-1 and MO1. Chimeras exhibiting notable phenotypic changes are highlighted in red. (**K**) Sequence logo plot displaying the polymorphisms across APN orthologs. Key secondary structures and critical residues involved in host range determination were marked with red dots. APN expression (red signal in F, I, and J) was validated via immunofluorescence targeting the C-terminal FLAG tag. Data are presented as mean ± SD for D. N-319 glycosylation sequon is labeled with Ψ in panels H and K. Statistical significance was determined by unpaired two-tailed *t* tests (n = 3 biological replicates).*: *P*<0.05, **: *P*<0.01, ***: *P*<0.005, and ****: *P*<0.001, NS: not significant. Scale bars: 100 μm for panels B, C, F, J, and I. See also Figure S3.

Both the European (RU15-1) and Asian (MO1) EriCoV strains bound efficiently to hedgehog APN (*E.amu* APN) (Figures 3B and S3B). In contrast, APNs from *Sorex araneus* (*S.ara*) and *Suncus etruscus* (*S.etr*), which are also members of the order Eulipotyphla, did not support detectable binding. Notably, two feline APN orthologs from the leopard (*Panthera pardus, P.par*) and domestic cat (*Felis catus*, *F.cat*) supported detectable RU15-1 and GER216 RBD binding, but not to MO1, indicating a strain-specific receptor compatibility (Figures 3B and S3B-3C). The receptor function of *P.par* and *F.cat* APNs was additionally validated across multiple EriCoV strains, and their ability to promote entry of GER216, FR7137, and PLJ64 highlighted the spillover potential of selected European EriCoV strains (Figures 3C-3D).

To define the molecular basis of this restricted tropism, we constructed domain-swapped chimeras between human and *E.amu* APNs and assessed their ability to support EriCoV RBD binding (Figure 3E). Substituting domain 2 of *E.amu* APN with its human counterpart abolished binding, while substituting *E.amu* Domain 2 with human APN counterpart restored binding, identifying domain 2 as a key determinant of species specificity. In contrast, swapping domains 1, 3, or 4 had minimal or no effect (Figure 3F). Flow cytometry further confirmed the key contribution of domain 2 to species specificity (Figure 3G). Fine mapping of domain 2 narrowed down the key host range determinants within region 288–331 (hAPN numbering), encompassing β16, β17, α1, and α2. Among them, β16 and α2 are more surface exposed, and the latter contains an N-319 glycosylation (Figures 3H-3J). Substitution of this region in hAPN with the *E.amu* counterpart (residues 279–321) enabled RBD binding (Figure S3D). Logoplot analysis based on the amino acid sequences of the 94 APN orthologs revealed high variability in this region, with *E.amu* APN displaying unique sequence features, particularly those around β16, as well as several key residues in α2 (Figure 3K). The partial sequence similarity of this region in *F.cat* and *P.par* APNs and the absence of an evolutionarily conserved N319-glycosylation, but not in human or *S.ara* orthologs, likely explains their receptor compatibility.

Collectively, these findings demonstrate a restricted APN tropism among EriCoVs and suggest that felids, including domestic cats, may serve as potential spillover hosts. Identification of host range determinants within the highly variable domain 2 highlights its central role in mediating EriCoV RBD interaction.

### Structure of *E.amu* APN in complex with RU15-1 RBD

To elucidate the molecular basis underlying *E.amu* APN recognition, we determined the cryo-EM structure of the RU15-1 RBD in complex with *E.amu* APN. 2D and 3D classification revealed that RU15-1 RBD binds exclusively to the monomeric form of *E.amu* APN, with no interaction observed with the dimeric form (Figures 4A and S4A). This observation is consistent with our results from size-exclusion chromatography and negative-stain electron microscopy (Figures S5A and S5B). Notably, the RU15-1 RBD–*E.amu* APN complex only accounted for approximately 5% of the total particles, which is consistent with the relatively low binding affinity and high dissociation rate of the complex determined by Bio-layer interferometry (BLI) assays (Figure S5C).

**Figure 4.**
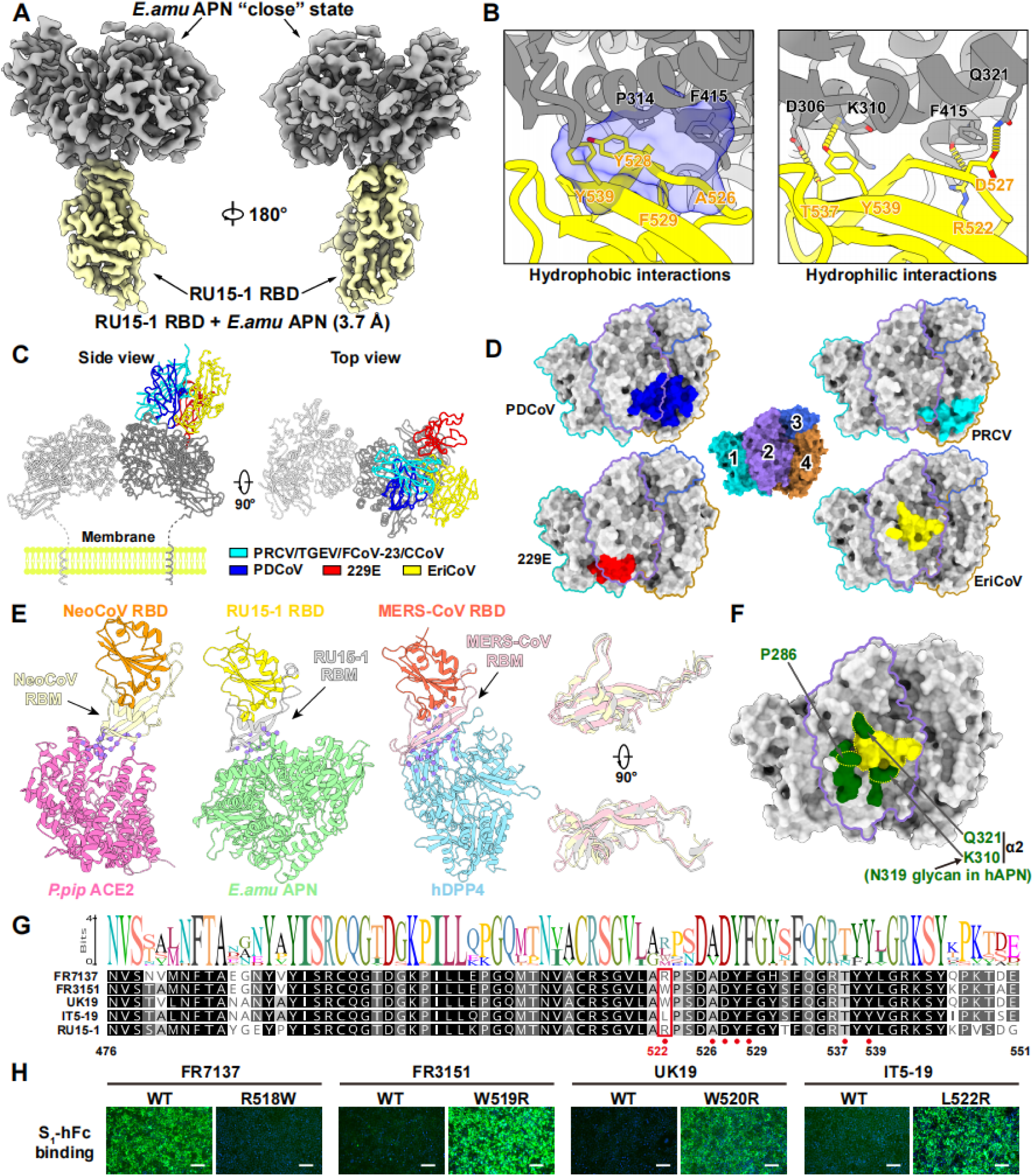
Cryo-EM structures of RU15-1 RBD and in a complex with *E.amu* APN. (**A**) The overall structure of the EriCoV RU15-1 RBD–*E.amu* APN complex. The RU15-1 RBD and *E.amu* APN are colored yellow and gray, respectively. (**B**) Interaction details between RU15-1 RBD and *E.amu* APN. Structures are shown as cartoon representations with key residues rendered as sticks. The hydrophobic pocket and hydrogen bonds are shown as a blue shadow with semi-transparency and yellow dashed lines, respectively. (**C**) Comparison of four distinct APN-binding modes used by the indicated coronaviruses, shown in both side and top views. PRCV (PDB: 4F5C), TGEV (PDB: 8Z27), FCoV-23 (9DAZ), and CCoV-HuPn-2018 (PDB: 7U0L) share a common binding mode, and only PRCV is shown as a representative. RBDs of PDCoV (PDB: 7VPP), HCoV-229E (PDB: 8WDE), and EriCoV (RU15-1) (this study) are shown in blue, red, and yellow, respectively. Only hAPN dimmer is shown for clarity, with individual monomers colored in silver and gray, respectively. (**D**) RBD footprints of four representative APN-using coronaviruses mapped onto APN surfaces. APN structures are color-coded: domain 1 in dark turquoise, domain 2 in medium purple, domain 3 in royal blue, and domain 4 in dark goldenrod. RBD footprints follow the same color scheme as panel C. (**E**) Structural comparison of three merbecovirus binding modes with distinct receptors: NeoCoV–*P.pip* ACE2 (PDB: 7WPO), EriCoV-*E.amu* APN (PDB: 9W6D) and MERS-CoV–hDPP4 (PDB: 4KR0). Right: Zoomed-in view showing the superimposed RBMs from each virus. All structures are shown in cartoon representation. The *P.pip* ACE2, *E.amu* APN, and hDPP4 are colored hot pink, light green, and sky blue, respectively. The RBDs of NeoCoV, EriCoV, and MERS-CoV are colored dark orange, gold, and tomato, respectively, with their corresponding RBMs highlighted in wheat, silver, and light pink. (**F**) Summary of the host range determinants (green) identified in this study and RU15-1 RBD binding interface (yellow dashed line) mapped onto *E.amu* APN surface. Residue positions in secondary structures are indicated; domain 2 is outlined in purple. (**G**) Sequence logo plot displaying the polymorphisms in receptor binding motifs from 22 EriCoV strains. Critical residues involved in APN interaction were marked with red dots. (**H**) Binding of wild-type and mutant S_1_-hFc proteins from the indicated European EriCoV strains to HEK293T cells stably expressing *E.amu* APN. Scale bars: 100 μm. See also Figures S4 and S5.

As expected, structural analysis of purified *E.amu* APN ectodomain proteins revealed that the monomer adopts two distinct conformations, designated as “closed” and “open” (Figures S4A, S5D, and S5E), reflecting the structural basis for APN’s multifunctionality.^71^ However, cryo-EM analysis demonstrated that RU15-1 binds exclusively to the “closed” conformation of *E.amu* APN (Figures 4A and S4A), indicating that RU15-1 binding is conformation-dependent, like PDCoV.^52^ In comparison, HCoV-229E can engage both the “open” and “closed” conformations of human APN, whereas PRCoV binds only to the “open” conformation of pig APN.^72,73^

We also determined the structure of the RU15-1 apo Spike. Cryo-EM analysis revealed that all RBDs adopt the “down” conformation (Figures S4A and S5F). This conformation is stabilized by an interaction network formed between glycans and neighboring protein surfaces: the N387 glycan on the RBD forms a hydrogen bond with N255 on the adjacent NTD, the N399 glycan on the RBD interacts with D527 on a neighboring RBD, and the N159 glycan on the NTD forms a hydrogen bond with S517 on the adjacent RBD. Collectively, these glycan–protein interactions lock the apo Spike in the “down” state (Figure S5G). A similar phenomenon has also been observed in other animal-derived apo spikes within the *Merbecovirus* subgenus, including HKU5, HKU25, and PDF-2180 (Figure S5H).

At the binding interface, the RBM of RU15-1 engages *E.amu* APN primarily through a combination of hydrophobic and polar interactions. Specifically, RU15-1 RBD residues Y539, Y528, F529, and A526 collectively form a hydrophobic pocket together with *E.amu* APN residues P314 and F415, contributing significantly to the binding affinity of the complex. In addition, Y539 and R522 on RU15-1 RBD form hydrogen bonds with backbone atoms of *E.amu* APN residues K310 and F415, respectively, thereby reinforcing local binding stability. Furthermore, T537 and D527 of RU15-1 establish additional hydrogen bonds with D306 and Q321 of *E.amu* APN, respectively, establishing an extensive interaction network that underpins specific recognition of *E.amu* APN by RU15-1 (Figure 4B).

Three distinct APN-binding modes have been reported among coronaviruses. Within the *Alphacoronavirus* genus, tegacoviruses TGEV, PRCV, CCoV-HuPn-2018, and FCoV-23 share a common binding mode, while HCoV-229E adopts a different one.^59,60,73–75^ The *deltacoronavirus* PDCoV^52^ also utilizes a distinct binding mode, featured by a slightly shifted binding orientation and footprint compared with tegacoviruses. (Figure 4C and S4B). Our cryo-EM reconstruction reveals that *betacoronavirus* EriCoV RBD binds to APN in a manner distinct from all previously known APN-using coronaviruses. Unlike the vertical binding orientations of tegacoviruses and

PDCoV, or the configuration of 229E, the EriCoV RBD binds APN in a laterally shifted orientation. Mapping the binding footprints of the four binding modes onto the APN structure showed that RU15-1 engages a previously uncharacterized region in the central portion of domain 2. Despite variations, all four RBDs engage domain 2, along with varying contributions from domains 1 and 4 (Figure 4D and S4B). This convergent usage of APN resembles the varying receptor-binding strategies observed for SARS-CoV-1, SARS-CoV-2, HCoV-NL63, NeoCoV, MOW15-22, and HKU5, all of which utilize ACE2 as their entry receptor.^40,41,50,51,54,57,76^ This EriCoV binding mode represents a novel mode of APN recognition by merbecoviruses, expanding our understanding of receptor usage across different coronavirus genera (Figure 4D).

Structural comparison of the EriCoV–*E.amu* APN complex with NeoCoV–*P.pip* ACE2 and MERS-CoV–hDPP4 complexes revealed that all three merbecoviruses utilize the same side of their RBM—primarily the β-sheet surface—to engage their respective receptors (Figure 4E). However, distinct indel patterns in the MERS-CoV RBM, as compared with the other two viruses, lead to conformational remodeling at the RBM tip, driving a shift in receptor specificity. Although EriCoV shares a similar RBM conformation with NeoCoV, multiple sequence differences at critical contact sites result in divergent receptor usage.

We then examined the molecular mechanisms of receptor species tropism from a structural perspective. In agreement with our functional data based on human and feline APN orthologs, all residues identified as critical for host range determination are situated in or around the RU15-1 RBD footprint, with some of them directly contributing to the RBD interaction, such as P286, K310 (corresponding to N319 in hAPN), and Q321 (Figure 4F).

To understand the impact of RBM polymorphism on receptor recognition, we generated a sequence logo plot of 22 EriCoV RBMs. Most residues critical for *E.amu* APN interaction are highly conserved (Figure 4G). However, the position of R522_RU15-1_ displays notable variability, featuring substitutions with distinct physicochemical properties. To explore the functional consequences of these variations, we generated several constructs with residue swaps at this site. Viral antigen binding assay revealed that R at this position confers higher binding affinity to *E.amu* APN compared to W and L, underscoring the role of EriCoV RBM polymorphism in modulating receptor binding affinity and probably ortholog specificity (Figure 4H).

### Hedgehog TMPRSS2-dependent spike activation

We next examined whether hedgehog APN could promote EriCoV entry and replication in human cells. As indicated by our earlier pseudovirus entry data (e.g., Figure 2J), EriCoV entry is highly dependent on trypsin treatment—a method commonly used to facilitate S protein activation via proteolytic cleavage.^77–79^ Analysis of S protein incorporation in selected EriCoV pseudovirus particles revealed inefficient proteolytic processing during particle biogenesis, similar to what has been reported for NeoCoV (Figure 5A).^40^

**Figure 5.**
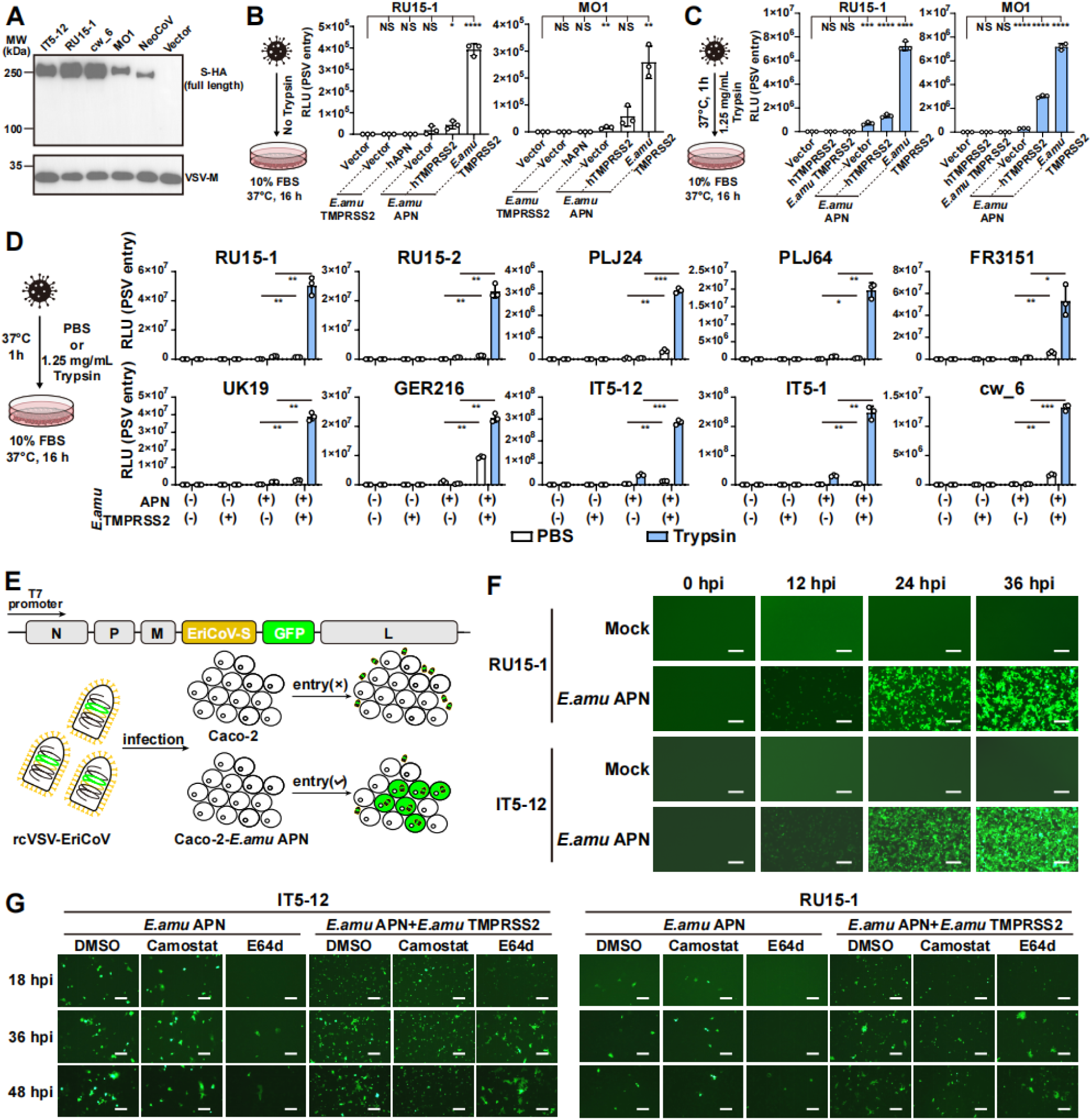
EriCoVs rely on hedgehog TMPRSS2 for efficient entry and propagation. (**A**) Detection of EriCoV S glycoprotein in VSV pseudovirions via Western blot using an anti-HA antibody targeting the C-terminally fused HA tag. VSV-M serves as a loading control. (**B-C**) Entry of RU15-1 and MO1 pseudovirus into HEK293T cells transiently expressing the indicated APN and TMPRSS2 orthologs. Flowcharts summarize the infection conditions with PBS (B) or trypsin (C) treatment. (**D**) Entry efficiency of selected EriCoV pseudoviruses into HEK293T cells expressing the *E.amu* APN, *E.amu* TMPRSS2, or both. PSVs were pretreated with 1.25 mg/mL Trypsin, and infection was conducted in the presence of 10% FBS without trypsin. (**E**) Schematic illustration of the genetic organization and amplification workflow for rcVSV-S constructs encoding EriCoV S glycoproteins. (**F**) Amplification of rcVSV-RU15-1 and rcVSV-IT5-15 in Caco-2 cells with or without the expression of *E.amu* APN. (**G**) Propagation of rcVSV-RU15-1 and rcVSV-IT5-12 in Caco-2 (left) or Caco-2-*E.amu* TMPRSS2 (right) cells with the transient expression of *E.amu* APN. Data are presented as mean ± SD for B, C, and D. Statistical significance was determined using unpaired two-tailed *t* tests (n = 3 biological replicates). *: *P*<0.05, **: *P*<0.01, ***: *P*<0.005, ****: *P*<0.001, and NS: not significant. Scale bars: 200 μm for panels F and G.

To investigate the role of host proteases in EriCoV entry and their cooperation with the receptor, we first assessed the impact of hTMPRSS2 or *E.amu* TMPRSS2 overexpression on pseudovirus entry without exogenous trypsin. Under this condition, pseudovirus entry of RU15-1 and MO1 was undetectable in HEK293T cells unless *E.amu* APN was co-expressed. Although the overall efficiency is low, entry was markedly increased upon co-expression of *E.amu* TMPRSS2, whereas hTMPRSS2 weakly enhanced entry, either with or without trypsin treatment (Figures 5B-5C). Notably, hAPN failed to support detectable entry even in the presence of *E.amu* TMPRSS2, confirming that hAPN is a poor receptor for EriCoV, and that *E.amu* TMPRSS2 functions in a receptor-dependent manner.

To determine whether this phenotype was specific to RU15-1 and MO1, we evaluated pseudoviruses of additional EriCoV strains. Since many strains showed lower infectivity, the pseudoviral particles were pre-treated with trypsin, followed by enzymatic inactivation before and during infection by 10% FBS. In HEK293T cells transiently expressing the different combinations of *E.amu* APN and *E.amu* TMPRSS2, all strains displayed similar dependence on APN and TMPRSS2 co-expression for more efficient entry (Figure 5D).

To further assess S-mediated viral propagation, we generated replication-competent VSV (rcVSV) encoding the S glycoproteins of IT5-12 and RU15-1 (Figure 5E).^40,50,51^ Viral amplification was observed in Caco-2 cells expressing *E.amu* APN without trypsin treatment (Figure 5F), but not in parental Caco-2 cells. In Caco-2 cells overexpressing *E.amu* APN only, rcVSV-RU15-1 and rcVSV-IT5-12 propagated over time and were sensitive to the cathepsin B/L inhibitor E64d, which indicates these viruses predominantly adopt the endocytosis pathway under this condition.^54,80^ Cells co-expression of *E.amu* APN and *E.amu* TMPRSS2 supported more efficient rcVSV amplification, and the boosted infection becomes less sensitive to E64d treatment, while syncytium formation reduced upon TMPRSS2 inhibitor Camostat treatment, ^54^ indicating *E.amu* TMPRSS2 allows more viruses to enter the cell through a cathepsin-independent cell surface fusion route (Figure 5G).

### Neutralizing antibodies against EriCoV

Next, we investigated whether RBD-targeting antibodies could block viral infection of HEK293T-*E.amu* APN cells. Mice were immunized with RBD proteins from the MO1 and RU15-1 strains, which share only 79% identity in their RBD sequences (Figure 6A). ELISA results confirmed that two immunizations generated specific antibodies against both the MO1 and RU15-1 RBDs, validating successful immunization (Figure 6B).

**Figure 6.**
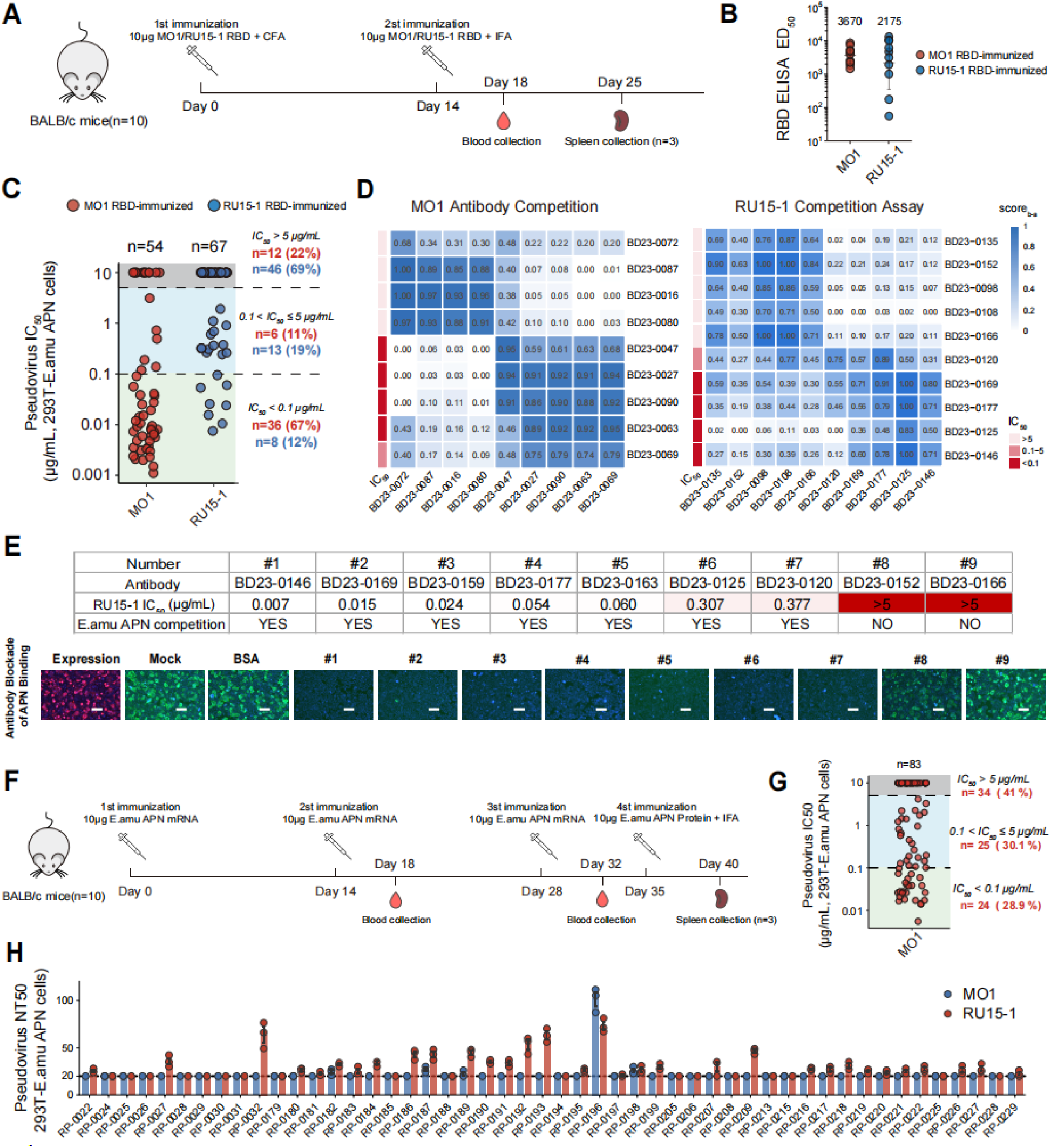
Neutralizing antibodies efficiently block EriCoV S-mediated entry. (**A**) Schematic of EriCoV RBD immunization in mice (n = 10 per group). (**B**) ELISA analyses of serum samples from immunized mice. ED_50_ values were calculated using four-parameter logistic regression. Geometric mean titers (GMT), fold changes, and statistical significance were determined by two-tailed *t* test and are shown above each group. (**C**) Pseudovirus IC_50_ values of monoclonal antibodies (mAbs) derived from MO1 (red circles, n = 54) or RU15-1 (blue circles, n = 67) immunization groups, categorized by potency. Neutralization assays were performed in HEK293T-*E.amu* APN cells using pseudoviruses corresponding to the respective immunogens. (**D**) Heatmap for pair-wise SPR competition scores of anti-RBD mAbs, with darker blue indicating stronger competition. Corresponding IC_50_ of mAbs are represented as red bars on the left of each heatmap, with darker red indicating higher neutralization potency. (**E**) Antibody blockage of APN binding of 9 representative mAbs isolated from RU15-1–immunized mice against RU15-1 pseudoviruses, determined using HEK293T-*E.amu* APN cells and RBD-hFc. (**F**) Schematic of *E.amu* APN immunization in mice (n = 10). (**G**) Pseudovirus IC_50_ values of mAbs derived from *E.amu* APN-immunized mice, categorized by potency. Neutralization assays were performed in HEK293T-*E.amu* APN cells using MO1 pseudoviruses. (**H**) Neutralization 50% titers of plasma samples from random human donors against MO1 and RU15-1 pseudoviruses, determined using HEK293T-*E.amu* APN cells. Dashed lines indicate the assay baseline. Scale bars: 100 μm for panel E. See also Figure S6.

Based on post-second-immunization ELISA titers, we selected the three highest-titer mice per group for splenocyte isolation. We sorted single B cells (7-AAD⁻/CD19⁺/IgM⁻/IgD⁻/OVA⁻/RBD⁺) and expressed monoclonal antibodies from them (Figure S6A). In total, we obtained 54 MO1 RBD-binding antibodies and 67 RU15-1 RBD-binding antibodies (Table S2). We evaluated these for their ability to neutralize MO1 and RU15-1 pseudoviruses in HEK293T-*E.amu* APN cells, which revealed potent neutralizing activity for some clones (Figure 6C). Among the 54 MO1-targeting antibodies, 36 (67%) exhibited strong neutralizing activity (IC_50_ < 0.1 μg/mL). In contrast, only 8 (12%) of the RU15-1-targeting antibodies displayed strong neutralizing activity, while 46 (69%) showed no neutralization.

We further validated antibody neutralization using replication-competent rcVSV-S pseudoviruses. Nine representative RU15-1 antibodies were tested against the RU15-1 and IT5-12 strains in Caco-2 cells expressing *E.amu* APN, alongside two well-characterized broadly neutralizing antibodies targeting the S2 domain as controls (Figure S6B).^81,82^ Strongly neutralizing antibodies potently inhibited replication of both strains, while weakly neutralizing antibodies showed only marginal suppression. These results aligned closely with IC_50_ values from single-round pseudovirus assays. Additionally, we found that all RU15-1 neutralizing mAbs could neutralize IT5-12 despite its RBD sharing only 87% sequence identity with RU15-1, suggesting conserved antigenicity among European EriCoV strains.

Next, we analyzed the relationship between the RBD-binding epitope and EriCoV neutralization activity of the antibodies. We performed epitope binning assays using SPR competition for antibodies binding to both the MO1 and RU15-1 RBDs (Figure 6D). The antibodies from both groups segregated into two distinct epitope bins, only one of which correlated strongly with pseudovirus neutralization. To determine if neutralization was mediated by blocking the receptor binding site, we selected nine representative RU15-1 antibodies for an APN receptor competition assay (Figure 6E). Recombinant RBD-hFc proteins were pre-incubated with RBD-binding antibodies and then applied to HEK293T cells transiently expressing *E.amu* APN to evaluate antibody APN blockage efficiency. The results revealed a clear correlation between the ability of antibody–APN blockage and viral neutralization (Figure 6E), underscoring the critical role of the RBD–APN interaction for EriCoV entry.

To further validate the role of APN as a functional receptor for EriCoV, we investigated whether antibodies targeting APN itself could block infection. Mice were immunized via intramuscular injection with 10 µg of *E.amu* APN mRNA on Days 0, 14, and 28. This was followed by a booster injection with 10 µg of purified *E.amu* APN protein formulated with incomplete Freund’s adjuvant (IFA) on Day 35 (Figure 6F). Mouse spleens were collected on Day 40, and *E.amu* APN-specific single B cells were sorted. Using the method described above, we expressed APN-binding antibodies from these cells. In total, we obtained 83 *E.amu* APN-binding antibodies (Table S2). Significantly, we found that over 50% of these antibodies displayed strong neutralization activity against EriCoV pseudoviruses in HEK293T-*E.amu* APN cells (Figure 6G). This result again provides direct evidence that APN is critical for EriCoV infection.

Together, we developed potent EriCoV-neutralizing antibodies that target the viral RBD–APN interaction and demonstrated their efficacy in blocking pseudovirus entry and propagation. These neutralizing antibodies represent promising candidate therapeutics for pandemic preparedness against potential future EriCoV spillover events. This is particularly relevant given that population immunity from SARS-CoV-2 infection and vaccination history shows negligible neutralizing activity against EriCoV (Figure 6H & Table S3).

## DISCUSSION

Coronaviruses circulating in wildlife continue to pose a significant threat of future pandemics.^83,84^ Predicting the cross-species transmission potential of specific coronavirus strains remains challenging due to limited knowledge of their host entry and transmission mechanisms. EriCoVs as relatives of MERS-CoV have been known for over a decade and are now recognized as genetically diverse and widely distributed.^25,33^ Unlike bats, which generally inhabit areas distant from human populations, hedgehogs often interact with humans as urban wildlife or pets.^85^ As important members of the merbecovirus subgenus associated with potential high pathogenicity, EriCoVs remain poorly characterized in terms of their spillover risks. A key obstacle to this understanding is the lack of receptor information, which hinders cross-species risk assessment, viral isolation, pathogenesis studies, and countermeasure development.

DPP4 was first identified in 2013 as a functional receptor for MERS-CoV^43^ and later considered the canonical receptor for merbecoviruses, especially after the identification of more DPP4-using HKU4-related viruses.^15,67^ However, many merbecoviruses appear unable to use DPP4 orthologs from either humans or their host species.^26,40,56,79^ A decade later, the identification of ACE2-using NeoCoV and others indicates that ACE2 is also an important receptor in mediating merbecovirus entry.^40,66^ Subsequent discoveries of two additional independently acquired interaction modes have established ACE2 as another major receptor in this subgenus.^24,50,51,55–57,86^ With most merbecovirus receptor usage now defined, EriCoV remained the last unresolved lineage. Notably, while EriCoV shows certain homology and similar insertion/deletion patterns with ACE2-using NeoCoV, these viruses do not appear to use ACE2.^40,56^ Some studies proposed that DPP4 may serve as an alternative receptor for EriCoVs through *in silico* protein docking simulations; however, no functional evidence supported this hypothesis.^33^

Our study identifies APN as its functional receptor, unexpectedly expanding APN usage to the betacoronavirus genus. APN was initially identified as a receptor for two alphacoronaviruses,^44,45^ each engaging it through distinct binding modes.^59,73,75^ More recently, deltacoronaviruses like PDCoV and avian CoVs were also shown to use APN via a third binding mode.^52^ Our findings add a fourth APN-binding mode, in which EriCoV binds exclusively to the ‘closed’ conformation of APN through a uniquely oriented RBD with a distinct footprint that evolved independently. This finding underscores the remarkable plasticity of coronavirus RBDs in adapting to diverse receptor surfaces^51^ and illustrates the limitations of predictive modeling, as even minor sequence changes can dramatically alter receptor usage of coronaviruses.^67^

The convergent utilization of APN by EriCoVs is reminiscent of the multiple independent acquisitions of ACE2 usage among coronaviruses.^50,51,86^ These observations suggest that the acquisition of a receptor by coronaviruses is not random. Intriguingly, the three merbecovirus receptors—ACE2, DPP4, and APN—are all ectopeptidases with abundant expression on epithelial surfaces of the respiratory and gastrointestinal tracts. The repeated exploitation suggests strong evolutionary advantages, such as more efficient entry, potential for broad tissue tropism, and favorable biochemical properties for virus–receptor interaction.^87^

Despite structural similarity and certain sequence homology between NeoCoV and EriCoV, their distinct receptor usage implies a receptor-switch event during their evolutionary separation. The direction and drivers of this shift remain unclear, though recombination, RBM insertions/deletions, and antigenic drift are likely contributors.^68,88,89^ Given their shared indel patterns, antigenic drift likely played a central role. In this case, adaptive mutations in multiple interacting residues are necessary to achieve entirely different receptor recognition modes,^58^ which aligns with their significant differences in receptor-interacting residues.

Our results show that EriCoVs are highly adapted to hedgehog APN orthologs, with limited ability to utilize APN orthologs from humans and other species, determined by critical residues and glycosylation near the interaction interface. The strong host specialization and narrow compatibility to other APN orthologs imply a high evolutionary barrier of these viruses to zoonotic transmission under current conditions. In addition to receptor binding, other host factors like TMPRSS2 compatibility, replication efficiency, and immune responses may also influence host range. Notably, EriCoV exhibits modest ability to engage feline APN, raising the possibility that domestic cats could serve as intermediate hosts. Given the close human–cat association, it will be important to monitor feline populations, especially in areas with known hedgehog–human overlap, for signs of EriCoV exposure. To date, no spillover events have been reported, and most known EriCoVs have been isolated from *E.eur* or *E.amu* hedgehogs. Whether these viruses infect other hedgehog species or have broader host ranges remains unclear.

Thus, continued surveillance and close monitoring of suspicious sequence changes, coupled with receptor-focused functional studies, are essential for pandemic preparedness.^11,83^ Although receptor-switch is not common for coronaviruses, merbecoviruses are notable for their RBM plasticity,^58,88,89^ facilitating their ability to adapt human receptors for zoonotic emergence. Development of EriCoV-specific neutralizing antibodies and vaccines could provide preemptive countermeasures. Overall, our findings expand the repertoire of receptor usage within the *merbecovirus* subgenus, providing a structural and functional basis for risk assessment and countermeasure development for those viruses that could potentially pose a threat to humans or domestic animals.

## Limitations of the study

Several limitations of this study should be acknowledged. First, all functional assessments of viral entry were conducted using pseudoviruses in cell-based systems, and no infection experiments with authentic viruses or *in vivo* models were performed. Moreover, due to the lack of available hedgehog-derived cell lines, we were also unable to assess viral entry in a native host cell context.

As such, our findings may not fully recapture the complexity of viral infection dynamics in natural hosts. Second, while we tested representative viral strains, the panel does not encompass the full genetic diversity within the lineage, and untested variants may exhibit distinct APN tropism features and protease preference. Lastly, although certain feline APN orthologs supported pseudovirus entry *in vitro*, it remains unclear whether the feline species are truly susceptible to infection *in vivo*. Receptor usage alone does not guarantee host susceptibility; other factors such as protease expression, immune barriers, and viral replication capacity also play critical roles. Future studies using authentic viruses, a broader range of viral strains, and *in vivo* challenge models will be essential to fully assess the zoonotic potential and pathogenicity of these merbecoviruses.

## Acknowledgments

This study was supported by the National Key R&D Program of China (2024YFC2607300 and 2023YFC2605500 to H.Y.), the National Natural Science Foundation of China (NSFC) projects (82322041, 32270164 to H.Y., 323B2006 to C.-B.M.), Scientific Research Innovation Capability Support Project for Young Faculty (ZYGXONJSKYCXNLZCXM-H16 to H.Y.), Natural Science Foundation of Hubei Province (2023AFA015 to H.Y.), the Fundamental Research Funds for the Central Universities (to H.Y.), and TaiKang Center for Life and Medical Sciences (to H.Y.). This work was also supported by the Strategic Priority Research Program (XDB1310000 to X.W.), National Science Foundation Grants (32325004, T2394482, and 12034006 to X.W.), Basic Research Program Based on Major Scientific Infrastructures, CAS-JZHKYPT-2021-05, CAS (YSBR-010 to X.W.), Ministry of Science and Technology of China (CPL-1233 to X.W.), and Changping Laboratory (2025D-04-01 to Y.C.). We thank Lu Lu (Fudan University) for providing EK1C4 peptides used in this study.

## Author contributions

H.Y., X.W., Y.C., and P.L. conceived the project. P.L., C.M., Z.M., Q.Z., J.L., and T.L. cloned S, RBD-hFc, and APN mutants and conducted RBD-hFc binding assays. C.M., P.L., J.L., Y.Y., T.L., and X.Yang designed APN ectodomain and antibody constructs and expressed recombinant proteins. C.L., P.L., and J.S. conducted phylogenetic and conservation analysis. Z.M., P.L., and M.H. conducted pseudovirus production and cell-cell fusion assays. Z.M., P.L., and X.Yang rescued the rcVSV-EriCoV-S pseudotypes and performed rcVSV propagation and inhibition assays. Q.Z. and T.L. conducted biolayer interferometry binding experiments. J.L., P.L., and X.Yang carried out VSV pseudovirus entry and neutralization assays. Q.Z. carried out cryo-EM sample preparation, data collection, and processing. Q.Z. and X.W. built and refined the structures. J.L., Y.Y., and Q.W. performed animal experiments and flow cytometric sorting of single B cells. Y.Y., J.L., and L.Y. conducted monoclonal antibody expression, functional characterization, and neutralization assays. P.L., Z. M., C.L., Y.S., J.S., X.Yu, J.L., and Y.X. analyzed the data. H.Y., X.W., and Y.C. wrote the manuscript with input from all authors.

## STAR ★ METHODS

### KEY RESOURCES TABLE

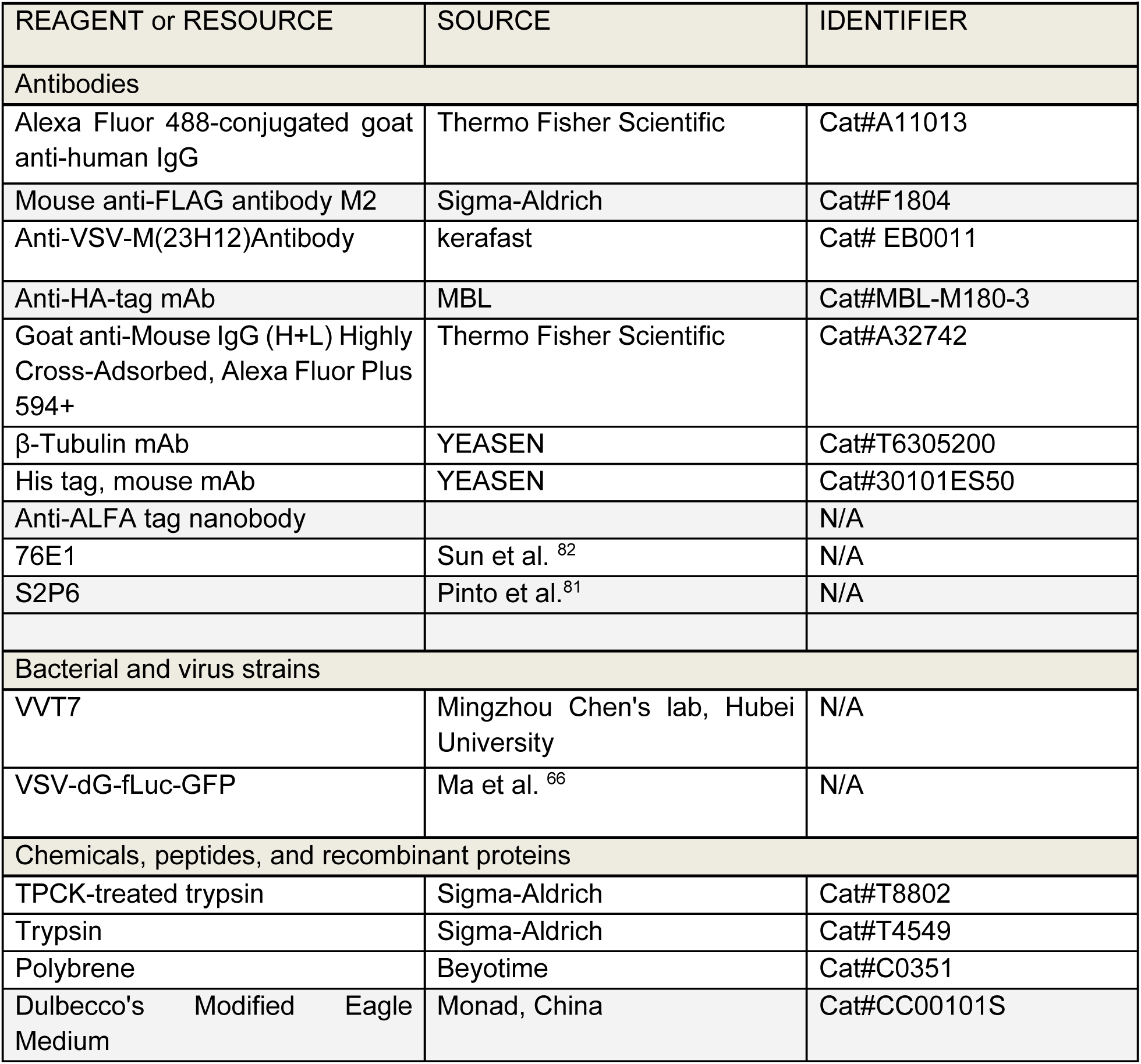
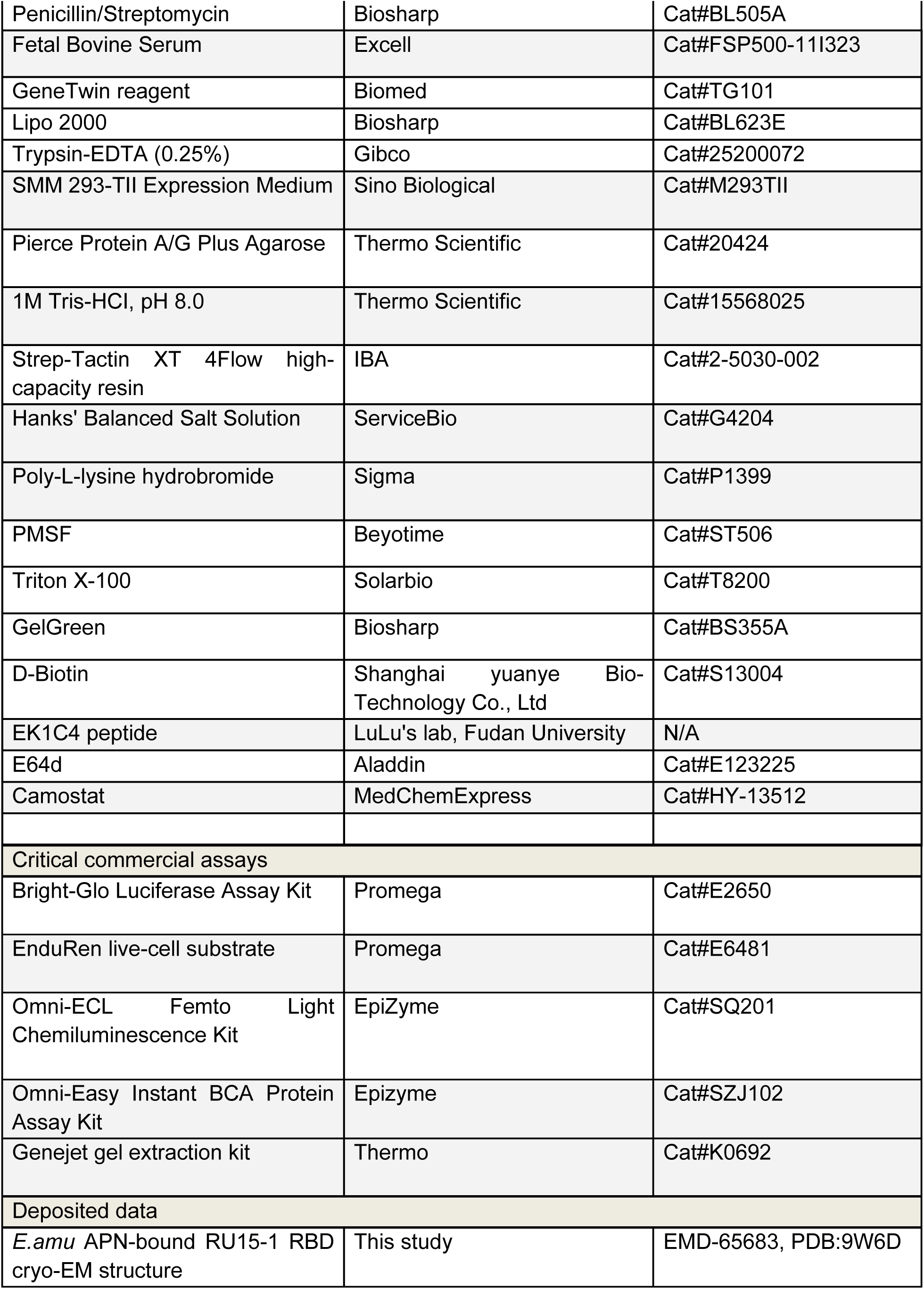
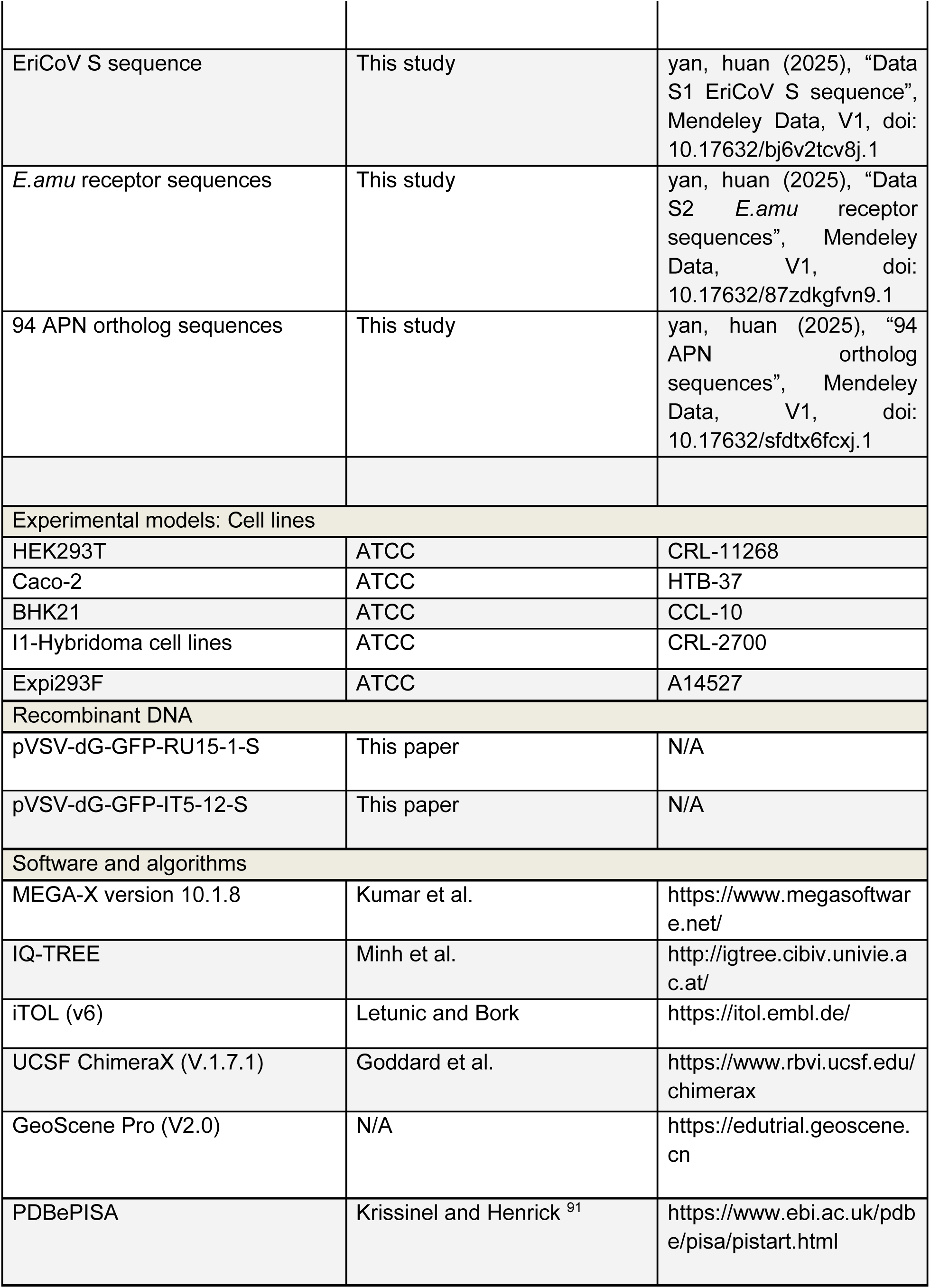
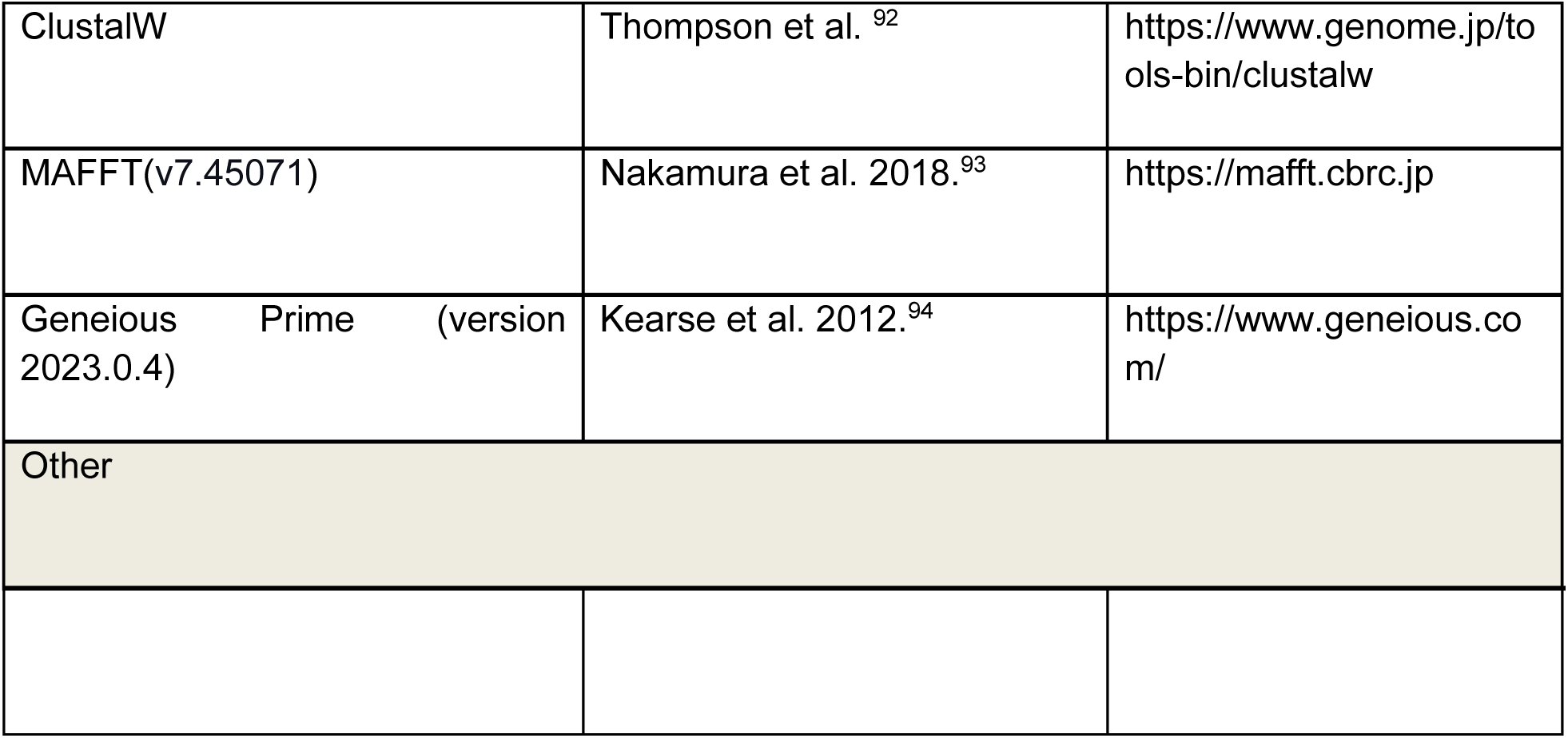

### RESOURCE AVAILABILITY

#### Lead contact

Further information and requests for resources and reagents should be directed to and will be fulfilled by the lead contact, Huan Yan (严欢)

#### Materials availability

All plasmids generated in this study are available with a completed Materials Transfer Agreement. Correspondence and requests for materials can be addressed to the lead contact.

#### Data and code availability

● The density map of the *E.amu* APN–RU15-1 RBD complex has been deposited in the Electron Microscopy Data Bank under the accession code of EMD-65683. The corresponding structural coordinate has been deposited in the Protein Data Bank: 9W6D.
● The 25 EriCoV spike sequences, four receptor sequences from amur hedgehog (*E.amu* ACE2, *E.amu* DPP4, *E.amu* APN, and *E.amu* TMPRSS2), and 94 APN-coding sequences have been deposited in Mendeley with DOIs and accession numbers listed in the key resources table.
● This paper does not report original code.
● Any additional information required to reanalyze the data reported in this paper is available from the lead contact upon request.

### EXPERIMENTAL MODEL AND STUDY PARTICIPANT DETAILS

### Cell lines and culture conditions

HEK293T (CRL-3216), Caco-2 (HTB-37), and I1-Hybridoma (CRL-2700) cells were obtained from the American Type Culture Collection (ATCC). HEK293T and Caco-2 cells were cultured in Dulbecco’s Modified Eagle Medium (DMEM, Monad) supplemented with 10% fetal bovine serum (FBS; Excell), 1% penicillin-streptomycin (PS; Thermo Fisher Scientific) at 37 °C in a humidified incubator with 5% CO₂. The I1-Hybridoma cell line, which produces a neutralizing antibody against vesicular stomatitis virus glycoprotein (VSV-G), was maintained in Minimum Essential Medium (MEM) with Earle’s balanced salts, 2.0 mM L-glutamine (Gibco), and 10% FBS. HEK293T or Caco-2 stable cell lines overexpressing various receptors were generated using lentivirus transduction, and then selected and maintained in growth medium with puromycin (1 μg/mL) or Zeocin (100 μg/mL).

#### Plasma donors

Plasma samples were collected from a random population at Beijing Ditan Hospital, Capital Medical University. All participants provided written informed consent prior to sample collection. The study was approved by the Ethics Committee of Beijing Ditan Hospital, Capital Medical University (Approval No. DTEC-KY2024-112-01). Demographic data for the plasma donors are presented in Supplementary Information, Table S3.

### METHOD DETAILS

#### Receptor sequence retrieval and assembly

The nucleic acid sequences of *E.amu* ACE2, *E.amu* DPP4, *E.amu* APN, and *E.amu* TMPRSS2 were assembled from RNA-seq raw data derived from *Erinaceus amurensis* Kidney (GSA accession ID: CRX555648) or *Erinaceus amurensis* Lymph (SRA Accession: SRX13651754). Briefly, raw next-generation sequencing (NGS) reads were downloaded from the Genome Sequence Archive (GSA) database and Sequence Read Archive (SRA) database and processed using Cutadapt (v2.4) to remove adapters and discard reads shorter than 30 base pairs. De novo assembly was performed with MEGAHIT (v1.2.9), using a minimum contig length of 100 bp. Resulting final contigs were analyzed using Geneious (2023.0.1) and were subjected to Geneious workflows for mapping reads to the indicated reference sequences under default parameters.

#### Plasmids and vectors

Plasmids for expressing coronavirus receptors or their derivatives were constructed by inserting human codon-optimized coding sequences obtained through commercial synthesis into the lentiviral transfer vector (pLVX-EF1a-Puro, Genewiz) with C-terminally fused 3×FLAG tags (DYKDHD-G-DYKDHD-I-DYKDDDDK).^40^ Human TMPRSS2 and *E.amu* TMPRSS2 expressing plasmid were constructed to pLVX-EF1a-Zeocin (clone based on pLVX-EF1a-Puro with the resistance gene replaced with Zeocin) with C-terminally fused ALFA-tag (PSRLEEELRRRLTE).^90^ DNA fragments for cloning chimera or mutants were generated by overlap extension PCR or gene synthesis and verified by commercial DNA sequencing.

For pseudovirus production, human codon-optimized spike sequences of MO1 (UMO75628.1), F6 (QGA70692.1), cw_6 (QOQ34381.1), IT8-1 (QRN68024.1), IT5-19 (QRN68031.1), IT5-17 (QRN68055.1), IT5-1 (QRN68048.1), IT5-11 (QRN68066.1), IT5-12 (QRN68078.1), IT5-15 (QRN68101.1), GER216 (AGX27810.1), GER174 (YP009513010), UK19 (QCC20713.1), PTH9 (XAJ09624.1), RU15-1 (XCN28775.1), RU15-2 (XCN28786.1), PLJ24 (XAJ09723.1), PLJ64 (XAJ09735.1), FR7137 (XNJ65105.1), FR3151 (XNJ65093.1), 229E (OQ920101), were cloned into the pCAGGS vector with 13-15 C-terminal residues replaced by a HA tag (YPYDVPDYA) to facilitate S incorporation.^95^

Plasmids expressing recombinant proteins, including coronavirus antigens, and soluble viral receptor proteins are described as follows. For CoVs S_1_/RBD-hFc fusion proteins, plasmids were constructed by inserting S_1_ coding sequences from MO1 (residues 18-734), F6 (residues 18-734), cw_6 (residues 18-734), IT8-1 (residues 20-735), IT5-19 (residues 20-737), IT5-17 (residues 20-737), IT5-1 (residues 20-735), IT5-11 (residues 20-740), IT5-12 (residues 20-734), IT5-15 (residues 20-737), GER216 (residues 20-736), GER174 (residues 20-736), UK19 (residues 20-734), PTH9 (residues 20-736), RU15-1 (residues 20-731), RU15-2 (residues 20-731), PLJ24 (residues 20-735), PLJ64 (residues 20-734), FR7137 (residues 20-731), FR3151 (residues 20-732); RBD coding sequences from RU15-1 (residues 366-574), MO1 (residues 366-575), PTH9 (residues 367-576), PLJ64 (residues 365-574), GER216 (residues 365-574), 229E (residues 293-434) into the pCAGGS vector containing an N-terminal CD5 secretion signal peptide (MPMGSLQPLATLYLLGMLVASVL) and C-terminal hFc-twin-strep-3×FLAG tags (WSHPQFEKGGGSGGGSGGSAWSHPQFEK-GGGRSDYKDHDGDYKDHDIDYKDDDDK) for purification and detection. For expressing soluble viral receptor proteins, plasmids were constructed by inserting ecto-domain coding sequences from *E.amu* APN (residues 60-960), human APN (residues 66-967) into the pCAGGS vector containing an N-terminal CD5 secretion signal peptide (MPMGSLQPLATLYLLGMLVASVL) and C-terminal 8×His tags for purification.

For Cryo-EM and BLI assays, DNA sequences encoding *E.amu* APN protein (residues 60–960; GenBank: XP_007516437.1) and RU15-1 RBD (residues 366–574; GenBank: XCN28775.1) were obtained from NCBI. An N-terminus CD5 signal peptide and a C-terminus His-tag were added to both constructs via PCR. These constructs were cloned into the pCAGGS mammalian expression vectors.

Plasmids for human APN gene knockout were constructed by inserting the single-guide RNA sequence into the pLentiCRISPRv3 vector (clone based on pLentiCRISPRv2-Puro with the resistance gene replaced with the mCherry coding sequence).

#### CRISPR-Cas9 knock-out cell line

Stable Caco-2 cell lines with knockout human aminopeptidase N (APN) were generated via the CRISPR/Cas9 system. Three guide RNAs (gRNAs) targeting the human APN gene were designed: gRNA1 (ATCGCCGTGGCCCGCCGCAA), gRNA2 (CGTTCAGGGCATAATCGCCG), and gRNA3 (CTTCTACCGCAGCGAGTACA). These guided RNAs were cloned into the lentiviral transfer vector pLVX-mCherry (Genewiz). Lentiviral particles were produced by co-transfecting HEK293T cells with the gRNA plasmids pMD2.G (Addgene #12259) and the packaging plasmids psPAX2 (Addgene #12260) at a 2:1:1 ratio using GeneTwin transfection reagent (Biomed, TG101). Viral supernatant was harvested to transduce Caco-2 cells. Single-cell clones were isolated by serial dilution at 72 hours post-transduction. APN knockout efficiency was confirmed by cDNA sequencing, 229E pseudotyped entry assays, and 229E spike protein-mediated cell-cell fusion assays.

#### Protein expression and purification

Proteins for binding, pull-down, or flow cytometry assays were produced in HEK293T cells via transient transfection, following the manufacturer’s guidelines. Supernatants were collected every 48 hours post-transfection. Twin-Strep-tagged proteins were enriched using Strep-Tactin XT 4Flow high-capacity resin (IBA, 2-5030-002), washed, and eluted with buffer BXT (100 mM Tris/HCl, pH 8.0, 150 mM NaCl, 1 mM EDTA, 50 mM biotin). Eluted proteins were concentrated, buffer-exchanged into PBS by ultrafiltration, aliquoted, and stored at −80 °C.

Protein expression and purification were conducted following the established method for Cryo-EM or BLI analysis.^96,97^ Briefly, the plasmids of these proteins were transiently transfected into suspension-grown HEK293F maintained at 37 °C with 8% CO_2_, rotating at 130 rpm. Supernatants were harvested 72 hours post-transfection, concentrated, and purified sequentially by Ni-NTA affinity chromatography and size-exclusion chromatography (SEC) using a Superdex 200 Increase 10/300 GL column (GE Healthcare) equilibrated with PBS buffer.

#### Enzyme-linked immunosorbent assay (ELISA)

ELISA plates were pre-coated with RBD protein from either strain MO1 or RU15-1 at a concentration of 1μg/mL in 0.05 M coating buffer (Solarbio, cat. no. C1055) and incubated overnight at 4 °C. After washing, plates were blocked, followed by the addition of 100 μL serially diluted mouse serum to each well and incubation at room temperature for 30 minutes. Plates were then washed and incubated with peroxidase-conjugated AffiniPure goat anti-human IgG (H+L) (Jackson ImmunoResearch, cat. no. 109-035-003) at 0.25 μg/mL for 30 minutes at room temperature. The reaction was developed using tetramethylbenzidine (TMB) substrate (Solarbio, cat. no. PR1200) and stopped by the addition of H_2_SO_4_. Absorbance was measured at 450 nm using a microplate reader (Thermo Scientific, Multiskan Fc).

#### Animal immunization and antibody screening

All animal experiments were conducted in accordance with protocols approved by the Animal Welfare Ethics Committee of HFK Biologics (Approval No.: HFK-AP-20250313).

Female BALB/c mice aged 6-8 weeks were used in the experiments. Mice were immunized according to the schedule outlined in Fig. 6A and F. Specifically, mice in the MO1 and RU15-1 RBD protein groups were immunized via subcutaneous injection with 10 µg of purified RBD protein from either the MO1 strain or RU15-1 strain, each formulated with Freund’s adjuvant. The primary immunization was administered with complete Freund’s adjuvant (CFA), followed by a booster injection formulated with incomplete Freund’s adjuvant (IFA) 14 days later. Blood samples were collected 4 days after the second immunization.

Mice in the *E.amu* APN group were immunized via intramuscular injection with 10 µg of *E.amu* APN mRNA on Day 0, Day 14, and Day 28, followed by a booster injection with 10 µg of purified *E.amu* APN protein formulated with incomplete Freund’s adjuvant (IFA) on Day 35. Blood samples were collected 4 days after the second immunization and 4 days after the third immunization. Serum titers were determined using enzyme-linked immunosorbent assay (ELISA).

Spleens from euthanized mice were minced in RPMI 1640 medium supplemented with 5% FBS, filtered through a 100 μm mesh, and treated with 1 × RBC Lysis Buffer (eBioscience, cat. no. 00-4333-57). After centrifugation (350 × g, 8 min), cells were resuspended in staining buffer (PBS +2% FBS) and adjusted to a final concentration of 1 × 10⁷ cells/mL. Recombinant biotinylated RBD proteins (MO1 or RU15-1 strain) or *E.amu* APN proteins were incubated with streptavidin-conjugated fluorophores (PE and APC; BioLegend) on ice for at least 30 minutes. Then H₂O was added to adjust volume, and the complexes were centrifuged at 14,000 × g for 10 minutes at 4 °C to remove aggregates. Before staining, splenocytes were incubated with TruStain FcX™ (BioLegend) for 10 minutes on ice in the dark to block Fc receptors. Antigen probes and fluorescently conjugated antibodies were then added, and cells were incubated on ice for 30 minutes in the dark. After staining, cells were washed twice with staining buffer (400 × g, 5–8 minutes) and resuspended in cold buffer. 7-AAD (eBioscience, cat. no. 00-6993-50) was added 10 minutes before flow cytometry to exclude dead cells. For cell sorting, antigen-specific B cells were defined as 7-AAD⁻, CD19⁺, IgM⁻/IgD⁻, OVA⁻, and either MO1 RBD⁺, RU15-1 RBD⁺, or *E.amu* APN⁺, depending on the immunization group. Single B cells were sorted using a MoFlo Astrios EQ Cell Sorter (Beckman Coulter) directly into 96-well PCR plates.

Sorted single B cells were lysed in buffer containing RNase inhibitor, 0.095% Triton X-100, Oligo dT primer, and dNTPs, followed by cDNA synthesis using the Vazyme R302 kit. Variable regions of heavy chains and light chains (lambda/kappa) were amplified via two-round PCR. After verification by 1% agarose gel electrophoresis and magnetic bead purification, the variable region amplicons were set aside. Separately, the human IgG1 constant region was cloned into pCMV3 vectors using homologous recombination with the ClonExpress II One Step Cloning Kit (Vazyme, cat. no. C112-02) at 37°C for 30 minutes. Variable regions were recombined into the pCMV3 vectors using the same homologous recombination method. Recombinant plasmids were transformed into DH5α competent cells (Tsingke, cat. no. TSC-C01), activated at 37°C, and plated for selection. Recombinant plasmids were transformed into DH5α competent cells (Tsingke, cat. no. TSC-C01), activated at 37°C, and plated for selection. Plasmid DNA of expanded cultures was extracted (CWBIO CW2105) after verified by the alignment results of Sanger sequencing. Protein production was carried out in Expi293F cells. Co-transfection was performed by mixing the antibody plasmids with polyethylenimine (Yeasen, 40816ES03) in 0.9% NaCl solution before adding to the cell culture. Cells were maintained under standard culture conditions. After 24 hours, 1 mL of nutrient supplement (OPM Biosciences, F081918-001) was added per culture bottle, with subsequent feedings every 48 hours during a 6–10 day culturing period. For antibody purification, supernatant containing the monoclonal antibodies was obtained by centrifugation (3,000 × g, 10 minutes). After incubation of Protein A magnetic beads (GenScript, L00695) with the supernatant for 2 hours, automated purification was achieved by using a KingFisher instrument (Thermo Fisher). Antibody concentration was measured with a NanoDrop (Thermo Fisher, 840-317400), while purity was assessed by SDS-PAGE (LabLead, P42015). Screening of monoclonal antibodies was performed using a pseudovirus neutralization assay. The antibody sequences are listed in Supplementary Information, Table S2

#### SPR competitive binding assay

Antibodies were assessed using the Biacore 8K system (GE Healthcare). For competitive binding experiments, His-tagged MO1 or RU15-1 RBD proteins were immobilized on Anti-His sensor chips (Cytiva) at a concentration of 1.35 μg/mL for 1 minute. Pairs of antibodies (37.5 μg/mL each) were then sequentially injected over the chip surface, with each injection lasting 2 minutes. The chip surface was regenerated with 1.5 M glycine. Competition between antibodies was quantified using the following score: score *_𝑏𝑏−𝑎𝑎_*=1−response*_𝑏𝑏_* / response^′^ where *response_𝑏𝑏_* represents the response units (RU) when antibody *b* was injected as the second antibody following antibody *a*, and *response^′^_𝑏𝑏_* represents the mean RU when antibody *b* was injected as the first antibody.

#### Antigen-hFc live-cell binding assay

Live-cell binding assays for coronavirus RBD-hFc fusion proteins were performed as previously described. HEK293T cells transiently expressing distinct receptors or orthologous APNs were incubated 48 hours post-transfection with recombinant RBD-hFc proteins (2 µg/mL diluted in DMEM) for 30 minutes at 37°C (5% CO₂). Cells were gently washed once with Hanks’ Balanced Salt Solution (HBSS), then stained for 1 hour at 37°C with Alexa Fluor 488-conjugated goat anti-human IgG (1 µg/mL; Thermo Fisher Scientific, #A11013) and Hoechst 33342 nuclear counterstain (1:10,000 dilution in HBSS). Both reagents were prepared in HBSS containing 1% bovine serum albumin (BSA). After staining, cells were washed twice with HBSS and imaged with fresh HBSS. Fluorescence microscopy used an MI52-N system (Nikon). Relative fluorescence intensities (RFU) were quantified using a Varioskan LUX multimode plate reader (Thermo Scientific).

For experiments with RBD-specific antibody competition, recombinant RBD-hFc proteins (2 µg/mL) and indicated RBD-specific antibodies (10 µg/mL) were mixed within 50 µL DMEM for 1 hour at 37°C and then applied to HEK293T cells transiently expressing *E.amu* APN for RBD-hFc binding efficiency evaluation.

#### Flow cytometry

To analyze RBD-hFc binding by flow cytometry, HEK293T cells transiently expressing the indicated APN orthologs were detached with 5 mM EDTA/PBS 24 hours post-transfection. Cells were incubated with RU15-1 or MO1 RBD-hFc proteins (2 µg/mL) at 4 °C for 1 hour. After two PBS washes, cells were stained for 1 hour at 4 °C using Alexa Fluor 488- or DyLight 594-conjugated goat anti-human IgG antibodies to detect RBD binding (Thermo Fisher Scientific; A11013). Following two additional PBS washes, 20,000 live cells (gated based on SSC/FSC) were analyzed on a CytoFLEX flow cytometer (Beckman).

#### Western blot

Western blotting was performed to assess S glycoprotein incorporation into VSV pseudoviruses. Pseudovirus-containing supernatants were concentrated through a 30% sucrose cushion (30% sucrose, 15 mM Tris-HCl, 100 mM NaCl, 0.5 mM EDTA) by centrifugation at 20,000 × g for 1 hour at 4°C. Pellets were resuspended in 1 × SDS loading buffer, boiled at 95°C for 30 minutes, and subjected to SDS-PAGE and immunoblotting.

To evaluate expression of different APN orthologs, HEK293T cells were transfected for 24 hours, lysed in 1% Triton X-100 at 4°C for 10 minutes, and clarified by centrifugation at 12,000 × g for 5 minutes. Supernatants were mixed with 5 × loading buffer and analyzed by Western blot. Membranes were blocked with 5% milk in PBST for 1 hour at room temperature, incubated overnight at 4°C with anti-HA or anti-APN polyclonal antibodies (Abclonal), and then probed with goat anti-mouse IgG secondary antibody for 1 hour at room temperature. After four washes in

PBST, signals were visualized using the Omni-ECL Femto Light Chemiluminescence Kit (EpiZyme, SQ201) and imaged on a ChemiDoc MP system (Bio-Rad).

#### Immunofluorescence assay

Immunofluorescence assays were performed to assess the expression levels of APN orthologs with C-terminal fused 3×FLAG tags. Cells expressing the receptors were seeded in the 96-well plates. Specifically, the transfected cells were fixed and permeabilized with 100% methanol for 10 minutes at room temperature. Then the cells were incubated with a mouse antibody M2 (Sigma-Aldrich, F1804) diluted in HBSS/1% BSA for 1 hour at 37 °C. After one HBSS wash, the cells were incubated with Alexa Fluor 594-conjugated goat anti-mouse IgG (Thermo Fisher Scientific, A32742) secondary antibody diluted in 1% BSA/PBS for one hour at 37 °C. The images were captured with a fluorescence microscope (Mshot, MI52-N) after the nucleus was stained blue with Hoechst 33342 reagent (1:2,000 dilution in HBSS).

#### Pull-down assay

For pull-down assays, recombinant viral RBD-hFc-Strep-FLAG and soluble *E.amu* APN-His proteins were expressed in eukaryotic cells and purified by affinity chromatography. Viral RBD-hFc-Strep-FLAG protein (1 μg/mL) was incubated with 30 μL suspended Mag-Strep beads in 500 μL PBS at 4 °C for 1 hour. Beads were washed three times with cold PBS and then incubated with 2 μg of *E.amu* APN-His protein in 1 mL PBS at 4 °C for an additional hour. After washing three times with cold PBS, beads were resuspended in PBS, denatured in 1 × SDS-loading buffer, and boiled for 20 minutes. Samples were analyzed by SDS-PAGE followed by western blotting.

#### Biolayer interferometry (BLI) binding assay

Binding affinities were measured using bio-layer interferometry (BLI) on an Octet RED96 instrument (Molecular Devices), following a protocol similar to that described above.^98,99^ Briefly, recombinant MO1 and RU15-1 RBD-His proteins (50 μg/mL) were immobilized onto Ni-NTA biosensors (ForteBio) for 300 s. Following loading, the biosensors were transferred to kinetics buffer (PBS supplemented with 0.02% Tween-20; PBST) for 60 s to remove unbound proteins. The sensors were then immersed in PBST containing serial dilutions (0–10 μM) of soluble *E. amurensis* APN ectodomain (twin-Strep-tag) for 120 s to monitor the association phase, followed by a 120 s dissociation phase in PBST. Data acquisition and analysis were performed using Octet Acquisition 9.0 and Octet Analysis 9.0 software (ForteBio), respectively.

#### Cell-cell fusion assays

A cell-cell fusion assay based on dual-split proteins (DSPs) was performed in HEK293T or Caco-2 cells stably expressing *E.amu* APN. Two groups of cells were prepared to assess the S glycoprotein-mediated membrane fusion promoted by receptor interaction. Group A cells were transfected with plasmids expressing S glycoprotein and rLucN(1-155)-sfGFP1-7(1-157), while group B cells received plasmids encoding the S glycoprotein and sfGFP8–11(158–231)-rLuc(156–311). At 12 hours post-transfection, both groups of cells were trypsinized, mixed, and seeded into a 96-well plate at 8 × 10^4^ cells per well. At 24 h post-transfection, the cells were washed twice with DMEM and then incubated with DMEM containing the indicated concentrations of Trypsin, from porcine pancreas (Sigma-Aldrich, T4549) for 30 minutes at 37 °C. Trypsin activity was then quenched by replacing the medium with DMEM supplemented with 10% FBS and Hoechst 33342 (1:5,000) for nuclear staining. Syncytia formation with green fluorescence was assessed after 1.5 hours post-trypsin treatment, and images were captured either by a fluorescence microscope (MI52-N; Mshot) or an ELISPOT reader (S6 UNIVERSAL M2).

#### Pseudovirus production

Pseudoviruses carrying coronavirus S glycoproteins were generated using a vesicular stomatitis virus (VSV)-based pseudotyping system based on a previously published protocol with minor modifications.^50^ Briefly, HEK293T cells were transfected with plasmids encoding the S proteins using Lipofectamine 2000 (Biosharp, China). After 24 hours, the cells were infected with VSV-ΔG-fLuc-GFP (1 × 10^6^ TCID50/mL) in DMEM supplemented with 8 μg/mL polybrene and incubated for 2 hours at 37 °C on a shaker. The inoculum was then replaced with fresh DMEM or SMM 293-TII Expression Medium (Sino Biological, M293TII) containing an anti-VSV-G monoclonal antibody (I1, 1 μg/mL) to neutralize residual input virus. After another 24 hours, the supernatant containing pseudoviruses was harvested, clarified by centrifugation at 13,523 × g for 5 minutes at 4 °C, and stored at −80 °C until use.

#### Single-round pseudovirus entry and neutralization assay

Single-round EriCoV pseudovirus entry assays were performed in HEK293T or Caco-2 cells transiently or stably expressing various receptors. Cells were seeded in 96-well plates at 3 × 10⁴ cells/well and infected with pseudoviruses (1.5 × 10⁵ TCID₅₀ in 100 μL) in the presence of the I1 anti-VSV-G neutralizing antibody to minimize background.

For proteolytic activation, pseudoviruses were pretreated with 1.25 mg/mL trypsin (Sigma-Aldrich, T4549) for 1 hour at 37°C. In experiments shown in Figures 2B–C, 2J, 3D, and 5D, trypsin activity was quenched with 10% FBS. In experiments for Figure 2G (RU15-1 only) 125 μg/mL trypsin was maintained during inoculation in serum-free DMEM. For untreated controls, PBS was added instead. At 16–20 hours post-infection (hpi), intracellular luciferase activity was measured using the Bright-Glo Luciferase Assay System (Promega, E2620) on a Thermo Varioskan LUX, or GloMax 20/20 Luminometer (Promega).

Monoclonal antibodies or plasma samples were serially diluted in DMEM (Hyclone, cat. no. SH30243.01) and incubated with pseudovirus in 96-well plates at 37°C under 5% CO_2_ for 1 hour. Trypsin-digested HEK293T-*E.amu* APN cells were then seeded into the plates and cultured for 24 hours. After discarding half of the supernatant, reconstituted lyophilized Bright-Lite Luciferase Assay Substrate was mixed with Bright-Lite Luciferase Assay Buffer (Vazyme, cat. no. DD1209-03-AB) before being added to the wells for incubation in the dark. Luminescence intensity was measured using a microplate spectrophotometer (PerkinElmer, model HH3400). The 50% inhibitory concentration (IC_50_) and the 50% neutralization titer (NT50) were calculated via four-parameter logistic regression using PRISM software (version 9.0.1).

#### rcVSV-CoV amplification and neutralization assay

Experiments involving replication-competent VSV-S (rcVSV-S) were approved by the Biosafety Committee of the State Key Laboratory of Virology and Biosafety of Wuhan University and performed under Biosafety Level 2 (BSL-2) conditions. Plasmids for rescuing rcVSV-S expressing IT5-12 or RU15-1 spike (S) glycoproteins were constructed by substituting the firefly luciferase (fLuc) coding sequence with coronavirus S gene sequences in the pVSV-ΔG-fLuc-GFP vector backbone.^66,87^ In these recombinant viruses, the VSV-G gene was replaced with genomically encoded S glycoproteins, with a GFP reporter expression cassette for visualization. rcVSVs expressing IT5-12 or RU15-1 S glycoproteins were generated via reverse genetics following a modified published protocol. Briefly, BHK21 cells seeded at 80% confluence in 6-well plates were infected with recombinant vaccinia virus expressing T7 RNA polymerase (vvT7 provided by Mingzhou Chen’s laboratory, Hubei University) at a multiplicity of infection (MOI) of 5 for 45 minutes at 37°C. Following infection, cells were transfected with pVSV-ΔG-GFP-S plasmids and helper plasmids (mass ratio: 5:3:5:8:1 for pVSV-ΔG-GFP-S:pBS-N:pBS-P:pBS-G:pBS-L). Supernatants containing rcVSV-S (designated P0) were harvested 48 hours post-transfection, filtered (0.22 μm) to eliminate residual vvT7, and used to infect Caco-2 cells pre-transfected with VSV-G for production of passage 1 (P1) viruses. To generate passage 2 (P2) viral particles carrying IT5-12 or RU15-1 S glycoproteins, P1 viruses were used to infect Caco-2 cells stably expressing *E.amu* APN without ectopic VSV-G expression and in the presence of anti-VSV-G (I1-hybridoma supernatant).

For viral amplification assays, 3×10⁴ trypsinized Caco-2 cells expressing relevant APNs were seeded in 96-well plates and incubated with rcVSV-CoV (1 × 10⁴ TCID_50_/100 μL) in DMEM supplemented with 2% FBS, with or without TPCK-treated trypsin. Nuclei were stained with Hoechst 33342 (1:10,000 dilution in HBSS) for 30 minutes at 37°C. Fluorescence images were acquired at designated time points post-infection using an MI52-N fluorescence microscope (Nikon).

For antibody neutralization assays, *E.amu* APN-expressing Caco-2 cells were seeded in 96-well plates at a density of 5 × 10^4^ cells per well. Twelve hours later, rcVSV-CoV was incubated with the indicated concentrations of antibodies in DMEM for 1 hour at 37 °C, then added to the target cells in DMEM containing 2 % FBS. At the indicated time post-infection, nuclei were stained with Hoechst 33342 (1:10,000 dilution in HBSS) for 30 minutes at 37 °C, and fluorescence images were acquired using a fluorescence microscope (MI52-N).

#### Cryo-EM sample preparation and data collection

Cryo-EM samples were prepared by mixing the respective protein components—either *E.amu* APN, RU15-1 RBD—at 1:3 molar ratios, followed by a brief incubation at 4°C overnight, similar to the previous frozen sample preparation.^100^ Each complex was subjected to size-exclusion chromatography (SEC) to ensure sample homogeneity. The samples were applied to glow-discharged holey carbon-coated gold grids (C-flat, 300 mesh, 1.2/1.3, Protochips Inc.), blotted for 7 seconds without blotting force under 100% relative humidity, and plunge-frozen in liquid ethane using a Vitrobot (FEI).

Cryo-EM data were collected on an FEI Titan Krios transmission electron microscope operated at 300 kV. Movies were recorded using a K3 Summit direct electron detector in counting mode, with each movie consisting of 32 frames (0.2 s per frame) and a total electron dose of 60 e⁻/Å². Automated data acquisition was performed using SerialEM. For the *E.amu* APN–RU15-1 RBD complex, data were acquired at a nominal magnification of 29,000 ×, corresponding to a calibrated pixel size of 0.83 Å. The defocus range was set between −1.2 and −2.0 μm.

#### Cryo-EM data processing, model fitting, and refinement

A total of 10,621 micrographs were collected for the *E.amu* APN–RU15-1 RBD complex. Beam-induced motion correction was performed using MotionCor2, and the defocus values of each micrograph were estimated with Gctf.^101^ A total of 3,191,475 particles were automatically picked and extracted for reference-free 2D classification using cryoSPARC.^102^ Based on the 2D class averages, 1,548,480 particles were selected for ab initio reconstruction in cryoSPARC with no symmetry imposed, resulting in multiple initial models representing potential conformational states of the complex.

The best candidate model was then selected and subjected to non-uniform refinement in cryoSPARC to generate the final cryo-EM density maps. The final resolution of each structure was estimated using the gold-standard Fourier shell correlation (FSC) criterion (threshold = 0.143), and local resolution was evaluated using ResMap.^103^ A detailed summary of the image processing workflow is provided in Supplementary Information, Figure S4A, and the statistics are listed in Supplementary Information, Table S1.

The atomic model of *E.amu* APN–RU15-1 RBD complex was generated by fitting *E.amu* APN and RU15-1 (predicted by AlphaFold3) into the cryo-EM densities described above by Chimera,^104^ followed by manual adjustment and correction according to the protein sequences and densities in Coot,^105^ as well as real space refinement using Phenix.^106^ Details of the refinement statistics of the complexes are summarized in Supplementary Information, Table S1.

#### Bioinformatic and structural analysis

A dataset of 25 unique EriCoV S glycoproteins was gathered via BLAST searches of the NCBI database using the HKU31-F6 S protein as a reference sequence. After eliminating redundant sequences with 100% RBD amino acid identity and those containing ambiguous base calls, 22 unique RBD sequences were collected for further analysis. The RBD and S protein sequences used for phylogenetic analysis in Figure 1C and Figure S1B were aligned via MAFFT-DASH, while the RBM sequence alignments in Figure 1F were aligned via MAFFT (v7.45071) with manual adjustments to optimize indel positioning. Phylogenetic trees were constructed using IQ-TREE (version 2.0.6) under the Maximum Likelihood model with 1,000 bootstrap replicates. Evolutionary model selection was performed using ModelFinder Plus in IQ-TREE with Bayesian Information Criterion (BIC) optimization. The best-fit models implemented were: WAG+G4 model for RBDs, GTR+F+I+I+R4 model for the whole genomes, WAG+F+I+R3 model for spike proteins, and Q.plant+F+I+I+R6 model for the APN protein sequences. Pairwise sequence identities were calculated in Geneious Prime (version 2023.0.4) (https://www.geneious.com/) following MAFFT-DASH alignment. Experimentally resolved structures used for structural analysis in this study included: MERS-CoV RBD (4KQZ), HKU4-1 RBD (4QZV), HKU5-19s RBD (9D32), NeoCoV RBD (PDB: 7WPO), EriCoV-GER216 RBD (9JMG), MOW15-22 RBD (9C6O), HCoV-229E RBD-hAPN complex (6U7E), PDCoV RBD-pAPN complex (7VPQ), and PRCV RBD-pAPN complex (4F5C). Structural visualization and analysis were performed using ChimeraX (v1.7.1). A total of 94 orthologous APN protein sequences from mammals and *Gallus gallus* were collected from NCBI (either directly annotated or extracted from whole-genome assemblies) and UniProt. Multiple sequence alignment was performed using MAFFT (v7.45071),^93^ and phylogenetic trees were constructed with IQ-TREE using the Maximum Likelihood method with 1,000 bootstrap replicates.^107^ The resulting trees were visualized using iTOL (https://itol.embl.de/).

### QUANTIFICATION AND STATISTICAL ANALYSIS

Most experiments related to pseudovirus infection were conducted 2-3 times with 2-3 biological repeats; technical repeats are indicated in the legends. Representative results were shown. All data were presented by MEAN or MEAN ± SD. Unpaired two-tailed t-tests were conducted for all statistical analyses for two independent groups using GraphPad Prism 10. *P*<0.05 was considered significant. *: *P*<0.05, **: *P*<0.01, ***: *P*<0.005, and ****: *P*<0.001, NS: not significant. No data were excluded for data analysis.

---

**Figure S1.**
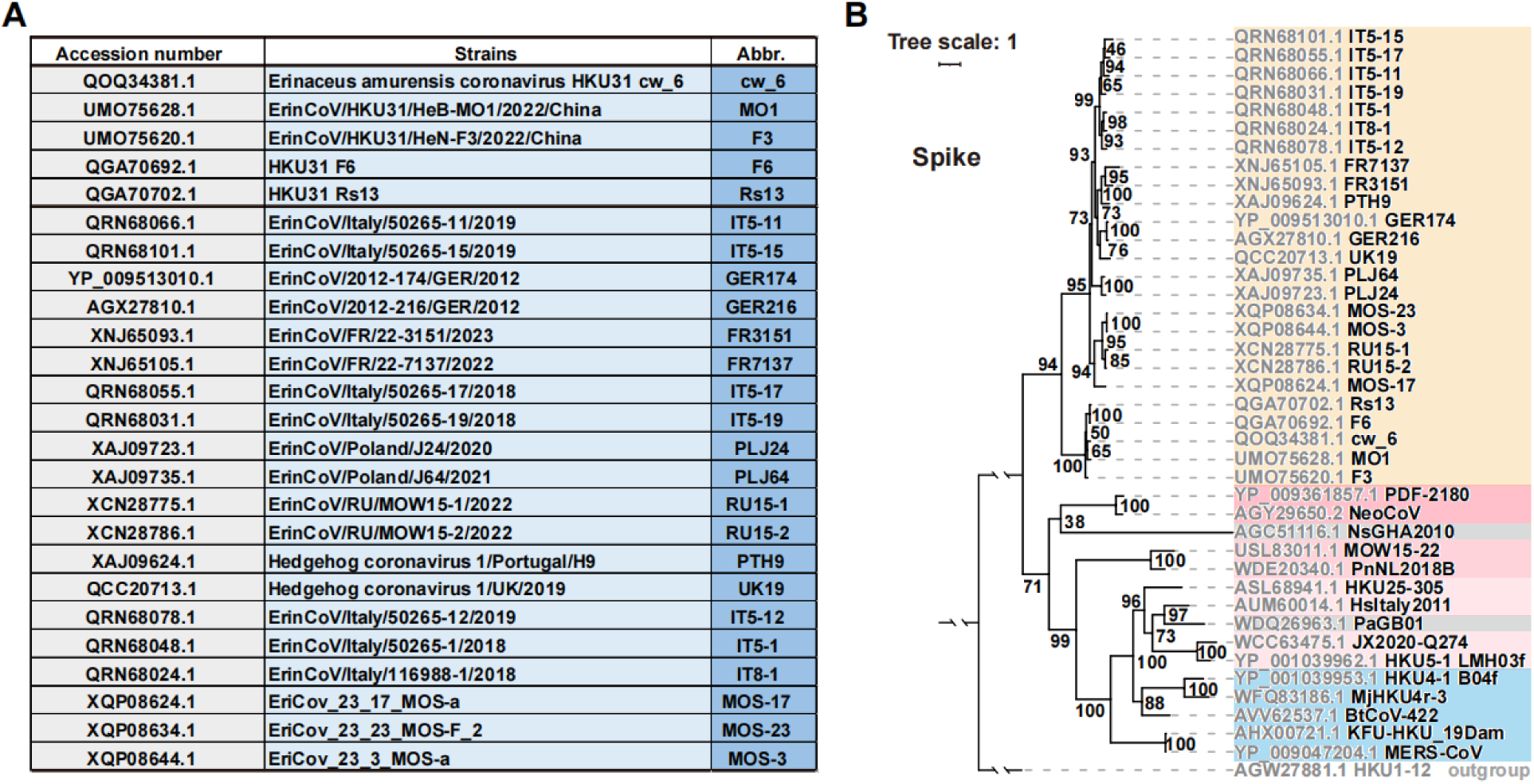
**Abbreviations and phylogenetic analysis of EriCoV S glycoprotein sequences.** (**A**) The accession number and abbreviation of the EriCoV strain names involved in this study. (**B**) Phylogenetic trees based on amino acid sequences of S glycoproteins from selected representative merbecoviruses were generated using IQ-TREE. Outgroup: HKU1-12.

**Figure S2.**
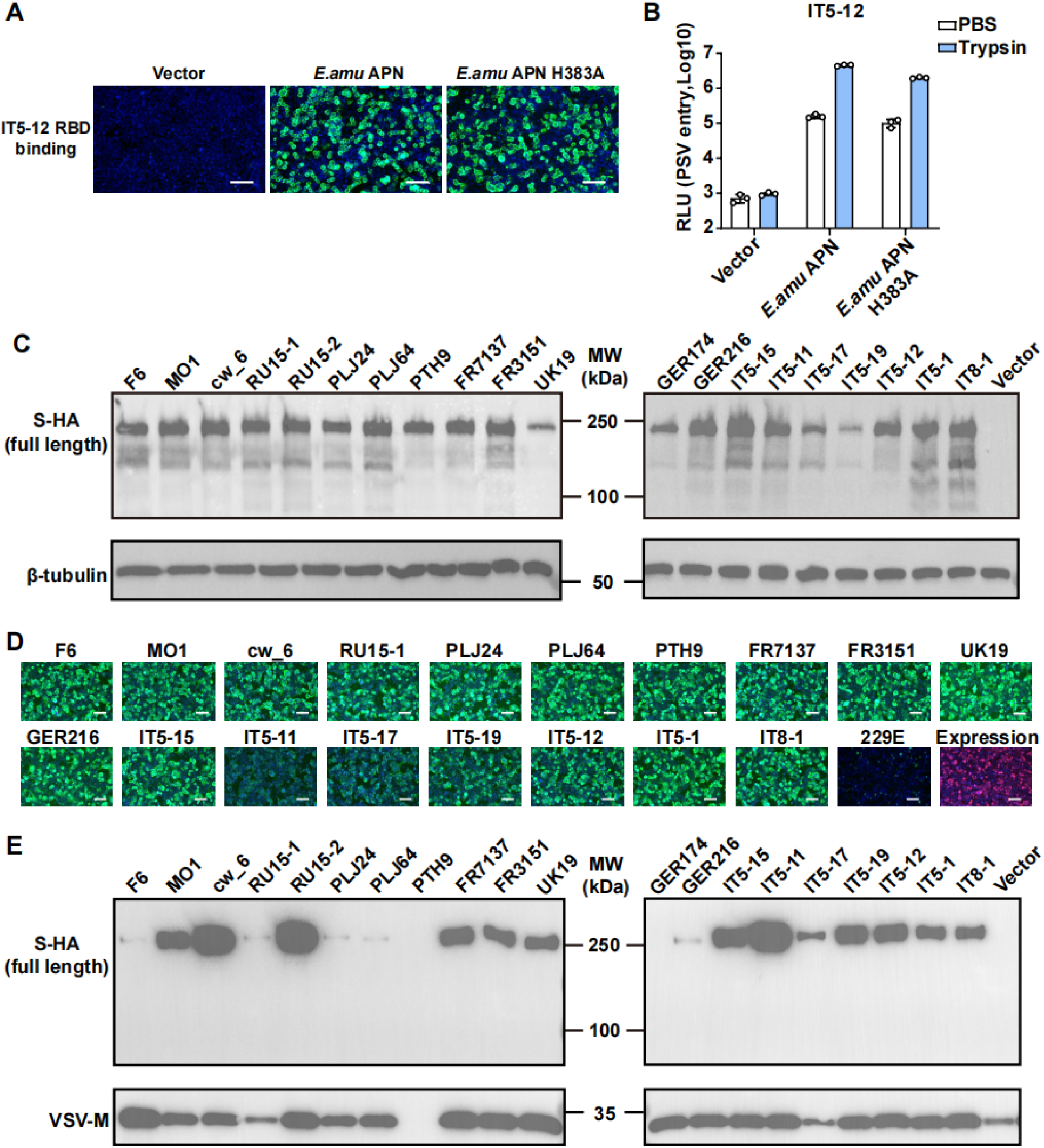
***E.amu* APN is utilized by various EriCoV strains in a catalytic activity-independent manner.** (**A**-**B**) *E.amu* APN receptor function is independent of its catalytic activity. IT5-12 RBD-hFc binding (A) and pseudovirus entry (B) in HEK293T cells transiently expressing WT *E.amu* APN or *E.amu* APN with H383A catalytic site mutation. Pseudoviruses were pretreated with 1.25 mg/mL trypsin. Mean ± SD for n=3 biological replicates. (**C**) Western blot analysis of indicated EriCoV S protein expression in HEK293T cells. β-tubulin served as a loading control. (**D**) EriCoV RBD-hFc binding in HEK293T cells transiently expressing *E.amu* APN, with 229E RBD as a negative control. (**E**) Detection of EriCoV S glycoprotein in VSV pseudovirions via Western blot using an anti-HA antibody targeting the C-terminally fused HA tag. VSV-M serves as a loading control. Scale bars: 100 μm for panels A and D.

**Figure S3.**
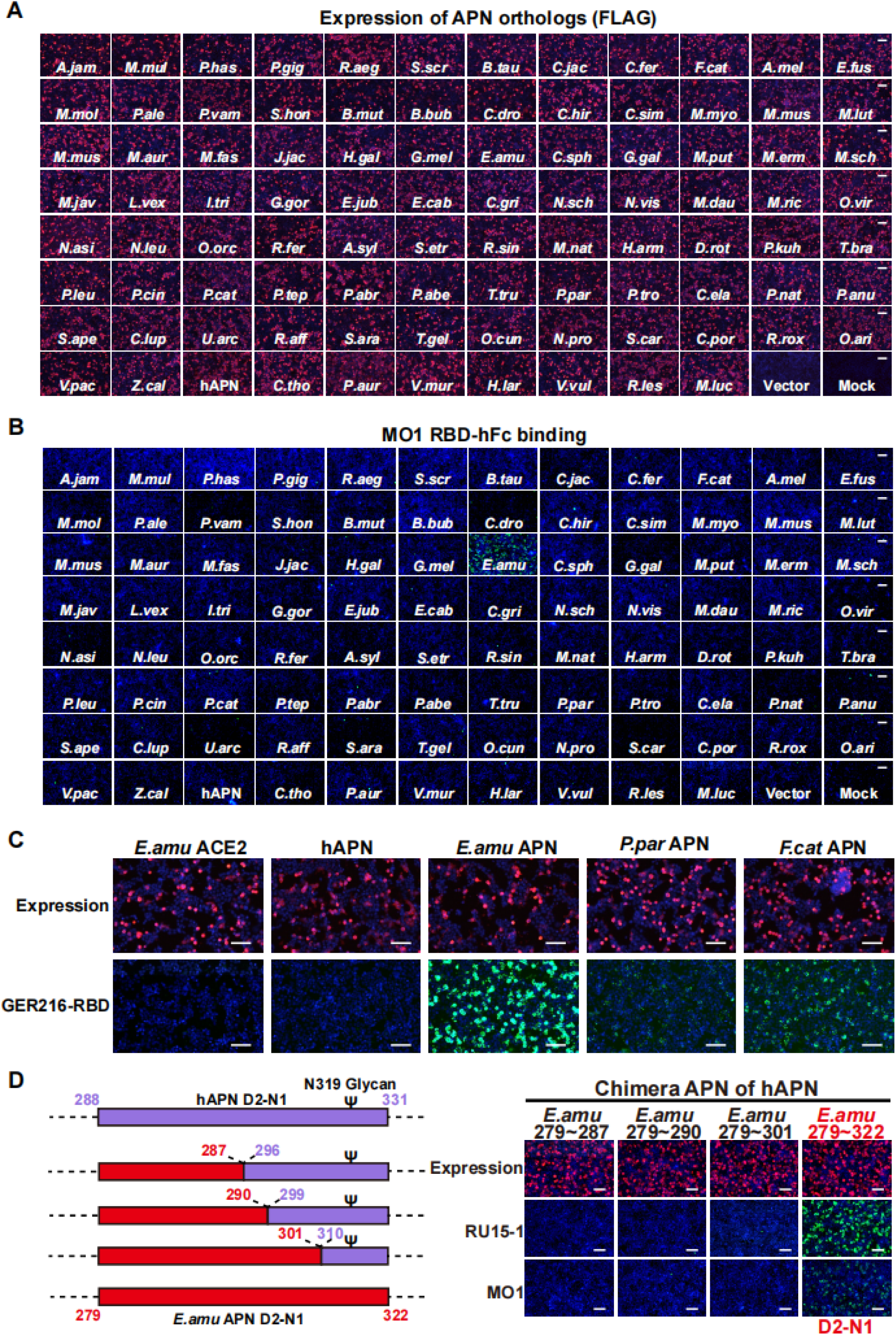
APN tropism and host range determinant of EriCoV. (**A**) Expression of 94 APN orthologs in HEK293T cells transiently transfected with C-terminally fused 3×FLAG tags was examined through immunofluorescence (red). (**B**) Binding of MO1 S_1_-hFc (green) to HEK293T cells transiently expressing the indicated APN orthologs, assessed by immunofluorescence using an anti-hFc antibody (green). (**C**) GER216 RBD-hFc binding (green) to HEK293T cells transiently expressing the indicated APN orthologs (red). (**D**) Substitution of the full D2-N1 region (residues 279-322) from *E.amu* APN is required for hAPN to support RBD-hFc binding of RU15-1 and MO1. N-319 glycosylation in hAPN is labeled with Ψ. Scale bars: 100 μm for panels A, B, C, and D.

**Figure S4.**
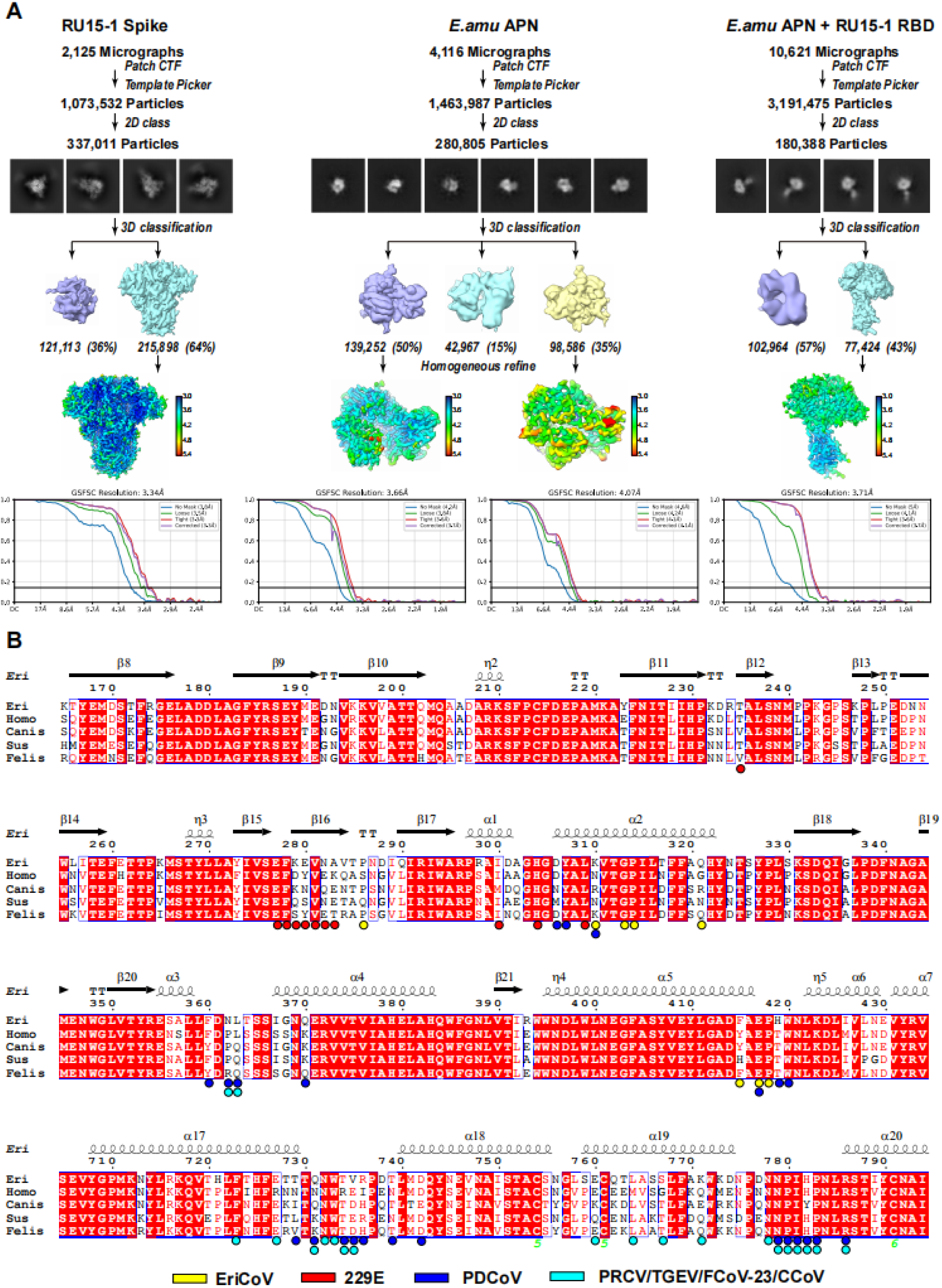
Cryo-EM data processing workflow and binding site analysis. (**A**) Flowcharts for structure determinations of the RU15-1 Spike, *E.amu* APN, and RU15-1 RBD–*E.amu* APN complex. Gold-standard Fourier shell correlation (FSC) curves are shown to assess map quality and resolution. Local resolution assessments of cryo-EM maps using rainbow are shown. (**B**) Multiple sequence alignment of APN orthologs from different species. APN sequences from *Erinaceus amurensis* (Eri), *Homo sapiens* (Homo), *Canis lupus* (Canis), *Sus scrofa* (Sus), and *Felis catus* (Felis) are aligned to highlight sequence conservation and variability. Binding sites located within 4 Å of viral RBDs are indicated by colored circles: yellow for EriCoV, red for HCoV-229E, blue for PDCoV, and cyan for PRCV, TGEV, FCoV-23, and CCoV, which share a common APN-binding mode.

**Figure S5.**
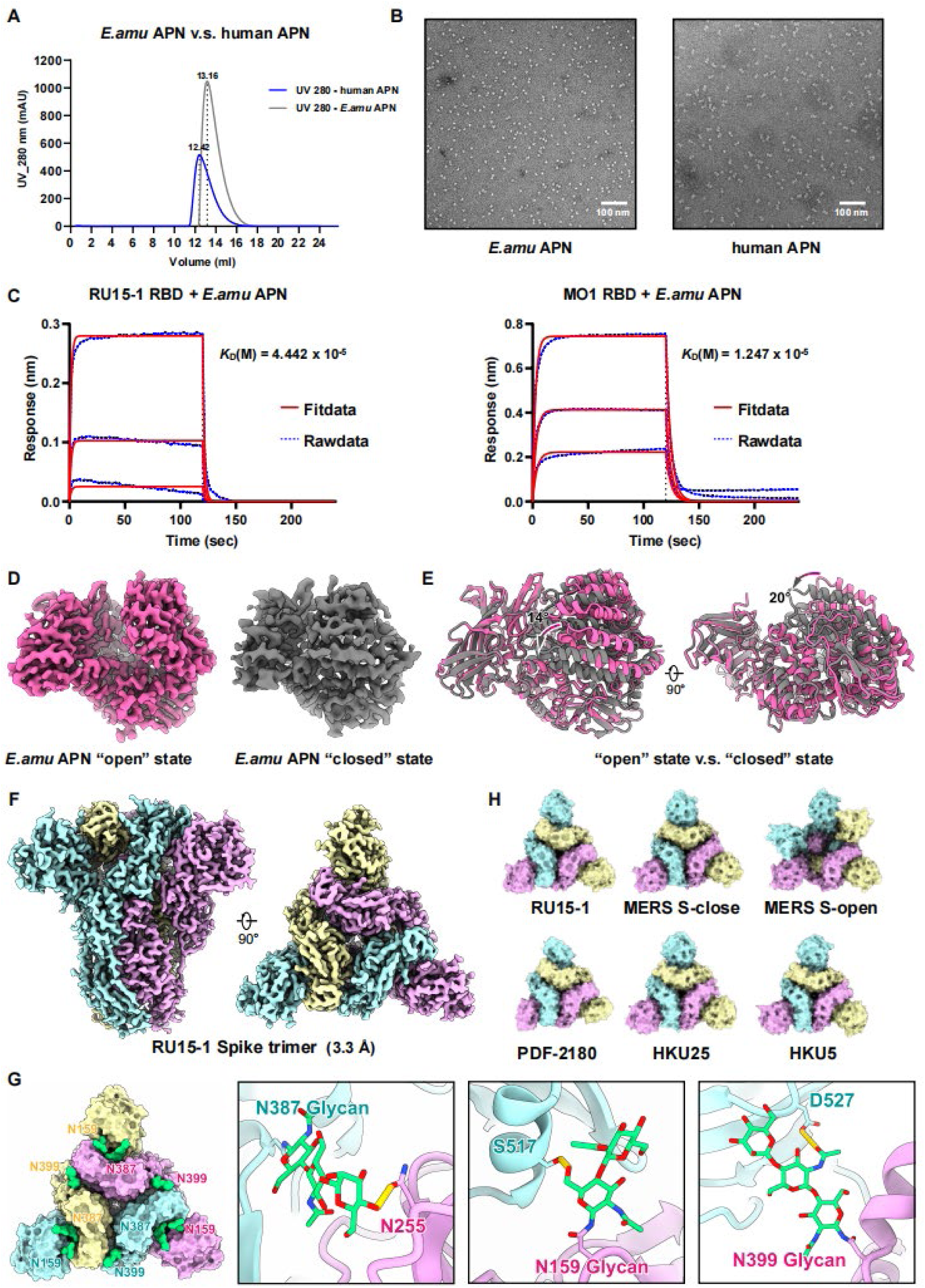
Conformations of *E.amu* APN, structural features of the RU15-1 spike and BLI assay. (**A**) Size-exclusion chromatography profiles of recombinant *E.amu* and human APN. Recombinant *E.amu* APN and human APN were expressed and purified using a Superdex 200 size-exclusion chromatography column. The elution profiles of size-exclusion chromatography traces are shown, with *E.amu* APN represented by the blue trace and human APN by the gray trace. (**B**)Negative-stain EM micrographs of *E.amu* APN (left) and human APN (right). (**C**) BLI assays analyzing the binding kinetics of *E.amu* APN ectodomains to immobilized monomeric RU15-1 RBD and MO1 RBD-His proteins on Ni-NTA biosensors. The BLI assay was performed using a 1:1 binding model with fixed-phase and mobile-phase interactions. Data were analyzed by global fitting, and the equilibrium dissociation constant (K_D_) was derived from the ratio of the association rate (ka) to the dissociation rate (kd). The fits are shown as red lines. (**D**) Conformational states of *E.amu* APN monomer. Cryo-EM structures of *E.amu* APN in the open (pink, PDB: 9ZA1) and closed (gray, PDB: 9ZA2) conformations are shown (left). (**E**) Structural superposition of the two conformations reveals a ∼14° angular displacement (middle left). Upon a 90° rotation, a larger deviation of ∼20° is observed between the open and closed states (middle right and right). (**F**) Cryo-EM structure of the RU15-1 spike trimer. Cryo-EM reconstruction of the RU15-1 spike trimer at 3.3 Å resolution shown in front (left) and top (right) views. The three protomers are colored cyan, purple, and yellow, respectively. (**G**) Structural representation of glycosylation sites on the RU15-1 Spike. N-linked glycosylation sites are shown in green (left: cartoon representation; right: stick representation). The three protomers are depicted in cyan, purple, and yellow, respectively, and key residues are annotated. (**H**) Cryo-EM reconstructions showing the conformational states of spike trimers from representative merbecoviruses, including RU15-1, MERS-CoV (open and close), PDF-2180, HKU25, and HKU5. The three protomers are colored cyan, purple, and yellow, respectively.

**Figure S6.**
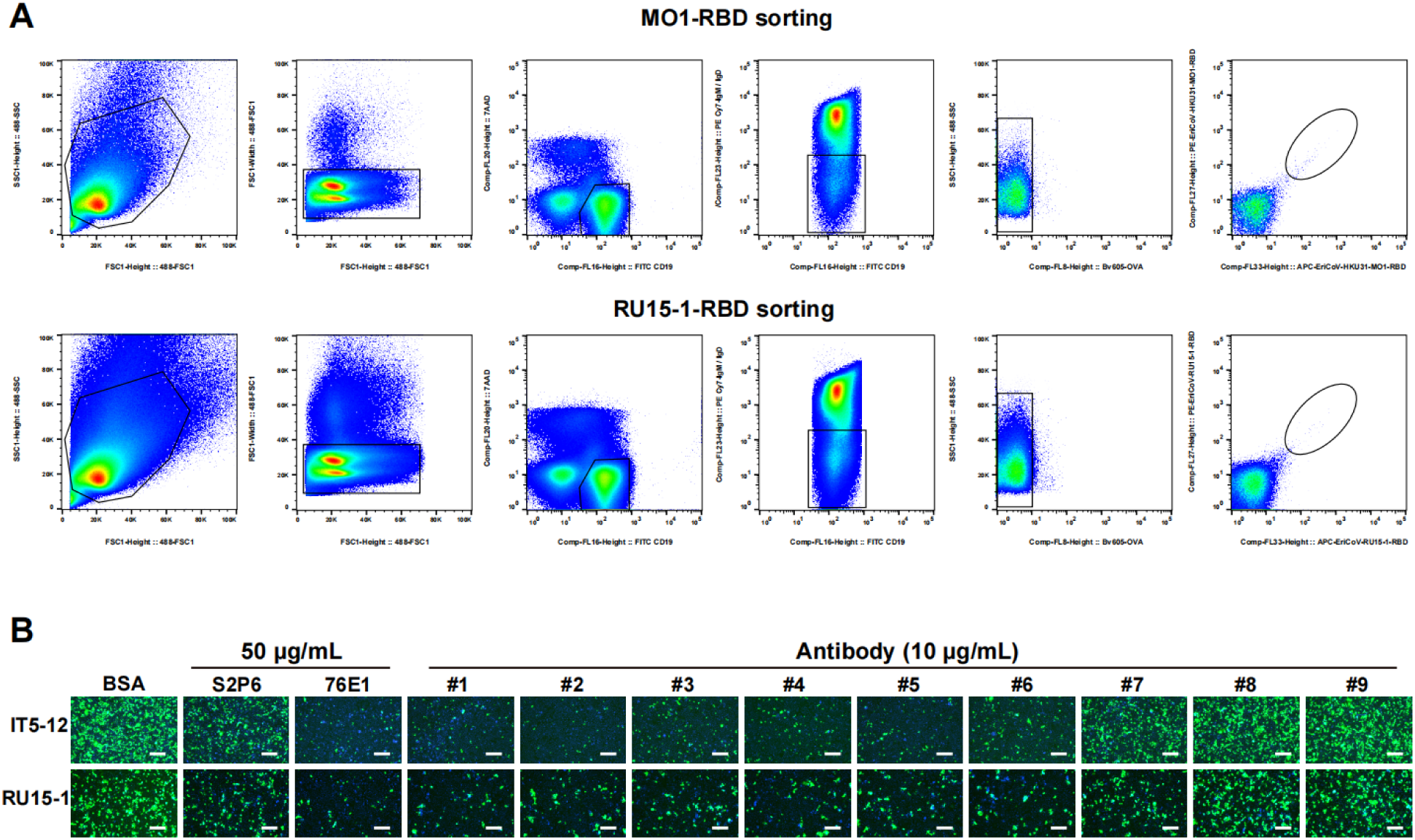
EriCoV RBD-specific antibodies efficiently inhibited the amplification of replication-competent VSV-EriCoV-S recombinant viruses. (**A**) Flow cytometry gating strategy for isolation of antigen-specific B cells. Splenic B cells were pooled from immunized BALB/c mice and stained for surface markers and antigen probes. The gating strategy is shown for two representative samples: MO1-RBD (top) and RU15-1-RBD (bottom). Lymphocytes were first identified based on FSC and SSC profiles, followed by doublet exclusion. Live cells were gated by excluding 7-AAD⁺ events. CD19⁺ B cells were selected, and IgM⁻/IgD⁻ class-switched B cells were further enriched. OVA-negative (BV605⁻) cells were gated to exclude non-specific binding. Antigen-specific B cells were identified as double-positive for PE-conjugated SA-MO1-RBD or SA-RU15-1-RBD and APC-conjugated SA-MO1-RBD or SA-RU15-1-RBD, respectively. These antigen-binding B cells were subsequently sorted for downstream single-cell V(D)J sequencing and antibody cloning. (**B**) Inhibition of rcVSV-EriCoV-S propagation by neutralizing antibodies using Caco-2 cells stably expressing the *E.amu* APN. IC_50_ values from single-round pseudovirus assays are shown above each panel. BSA: Bovine serum albumin, 10 μg/mL. Scale bars: 200 μm.

## Supplementary information for

**Table S1** Cryo-EM data collection and atomic models refinement statistics

**Table S2** Sequence information of 121 EriCoV-specific antibody sequences

**Table S3** Demographic data for the plasma donors

